# Beyond Static Brain Atlases: AI-Powered Open Databasing and Dynamic Mining of Brain-Wide Neuron Morphometry

**DOI:** 10.1101/2024.09.22.614319

**Authors:** Shengdian Jiang, Lijun Wang, Zhixi Yun, Hanbo Chen, Jianhua Yao, Hanchuan Peng

**Affiliations:** New Cornerstone Science Laboratory, Institute for Brain and Intelligence, Southeast University, Nanjing, China; Tencent AI, Shenzhen, China; Shanghai Academy of Natural Sciences, Shanghai, China

## Abstract

We introduce NeuroXiv (neuroxiv.org), a large-scale, AI-powered database that provides detailed 3D morphologies of individual neurons mapped to a standard brain atlas, designed to support a wide array of dynamic, interactive neuroscience applications. NeuroXiv offers a comprehensive collection of 175,149 atlas-oriented reconstructed morphologies of individual neurons derived from more than 518 mouse brains, classified into 292 distinct types and mapped into the Common Coordinate Framework Version 3 (CCFv3). Different from conventional static brain atlases that are often limited to data-browsing, NeuroXiv allows interactive analyses as well as uploading and databasing custom neuron morphologies, which are mapped to the brain atlas for objective comparisons. Powered by a cutting-edge AI engine (AIPOM), NeuroXiv enables dynamic, user-specific analysis and data mining. We specifically developed a mixture-of-experts algorithm to harness the capabilities of multiple large language models. We also developed a client program to achieve more than 10 times better performance compared to a typical server-side setup. We demonstrate NeuroXiv’s scalability, efficiency, flexibility, openness, and robustness through various applications.

## Main

Neuronal morphologies, characterized by their diverse branching patterns and anatomical arborization, provide critical insights into cell types and brain functional networks (Zeng & Sanes, 2017; Luo, 2021; Zeng, 2022). Recent advancements, including sparse labeling techniques (Aransay et al., 2015; Karube et al., 2004; Rotolo et al., 2008), high-resolution brain imaging (Economo et al., 2016; Gong et al., 2016; Zhong et al., 2021), terabyte-scale image handling (Bria et al., 2016; Peng et al., 2017; Y. Wang et al., 2019), and neuron tracing methods (Peng et al., 2010, 2011; Xiao & Peng, 2013; Feng et al., 2015; Jiang et al., 2022; Manubens-Gil et al., 2023), have significantly enhanced our capability to digitize brain-wide neuron morphologies. As a result, there has been a substantial increase in the volume of publicly accessible neuron morphologies, which supports various quantitative analyses, including the morphological characteristics of individual neurons (Peng et al., 2021; Winnubst et al., 2019), dendritic microenvironments (Y. Liu et al., 2023), neuron typing (L. Liu et al., 2023; Xiong et al., 2024), and the organizational principles of neuron projections (Gao et al., 2022, 2023; Jiao et al., 2023; Qiu et al., 2024). However, a remarkable gap exists: how to harness these valuable datasets from diverse sources for new knowledge discoveries while addressing dynamic needs throughout the development process (Wilkinson et al., 2016; Martone, 2024).

Current data dissemination solutions for neuron morphology (Akram et al., 2018; Winnubst et al., 2019; Kenney et al., 2024; Qiu et al., 2024) generally fall into two categories: browser-based atlases, such as Neuron Browser (mouselight.janelia.org) and Digital Brain (<u>mouse.digital-brain.cn</u>), and archiving platforms, including Brain Image Library (brainimagelibrary.org) and NeuroMorpho.Org (neuromorpho.org). Archiving platforms typically provide dataset downloads and offer a broader range of data from various sources, while browser platforms furnish additional data exploration tools—such as visualization and statistical analysis—but often limit access to data from their respective laboratories (**Extended Data Table 1 and Supplementary Note 2**). Large-scale offline analyses, which require aggregating datasets from multiple sources, introduce further neuroinformatics challenges. These include dataset harmonization, alignment with common coordinate frameworks (CCFs), extraction of key metadata such as morphological and anatomical features, and transforming neuronal features and patterns into meaningful insights. Addressing these challenges demands both domain-specific expertise and advanced coding skills (**Supplementary Note 1)**.

We introduce the NeuroXiv platform (neuroxiv.org), currently hosted on Amazon Web Services (AWS), designed to address challenges in databasing and mining brain-wide neuron morphometry. Building upon the foundational work of the Allen Brain Atlas (Q. Wang et al., 2020) and NeuroMorpho.Org, we have expanded efforts to establish a standardized atlas-oriented database of neuron morphometry (**Fig. 1A and Methods**). To address challenges associated with large-scale analysis of neuron morphology, we have developed the AI-Powered Open Mining (AIPOM) engine. This engine offers functionalities such as searching and visualizing neuron morphometry, enabling analyses including data statistics, cell typing, and connectivity studies. Crucially, it incorporates advanced capabilities, such as generating AI-driven mining reports (**Fig. 1A and Fig. 2A**).

**Fig. 1.**
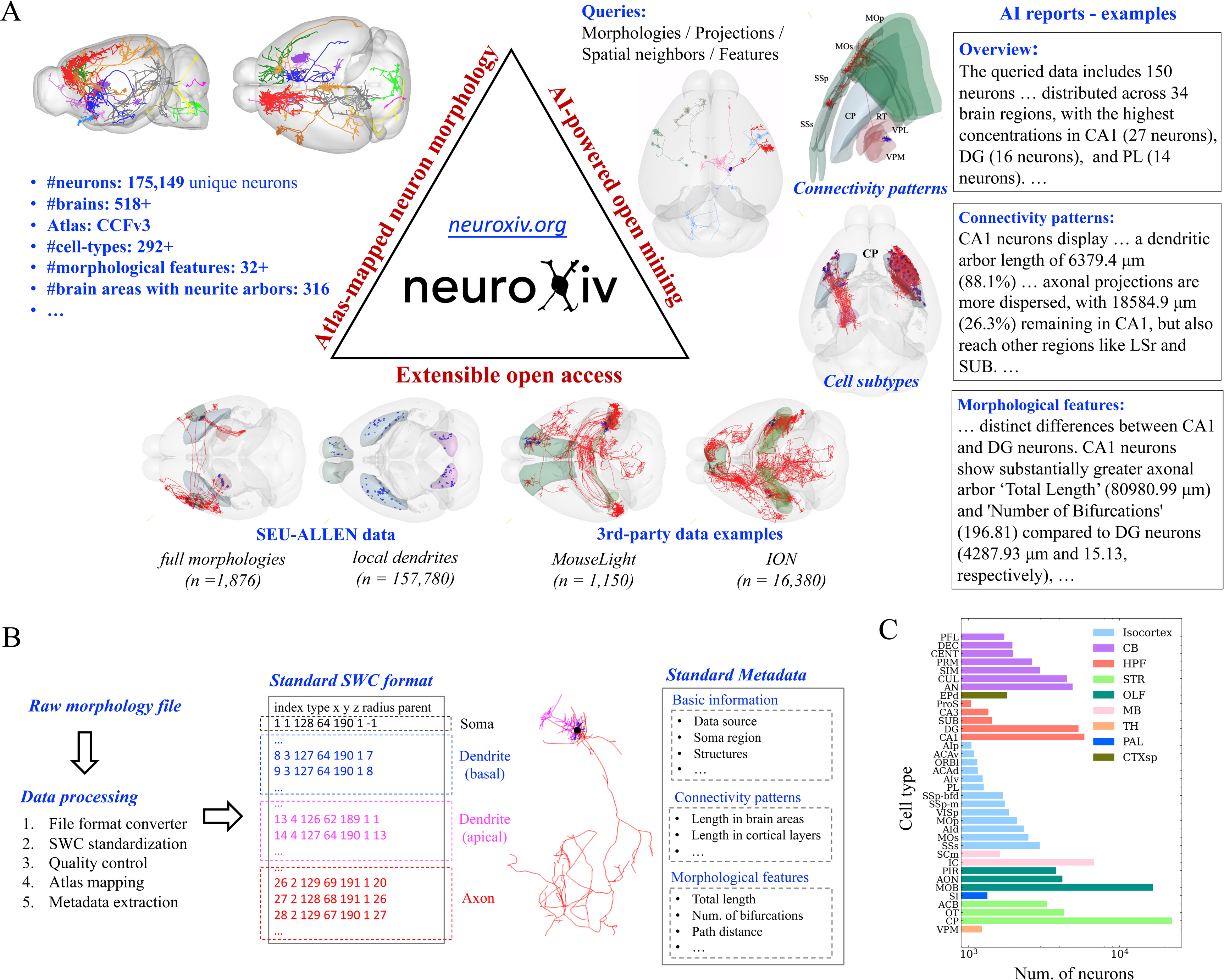
Overview of NeuroXiv, an open, AI-assisted database for interactive brain-wide neuron analysis. **A**, NeuroXiv is founded on three pillars: atlas-mapped neuron morphology, AI-powered open mining, and extensible open access. AIPOM engine enables users to efficiently and flexibly retrieve neuronal data, explore neuron types and connectivity patterns, and offers an intelligent mining tool for generating comprehensive data mining reports. **B,** the dataset standardization process in NeuroXiv is performed server-side, which is crucial for ensuring data reusability and interoperability. This process involves formatting raw morphology files into the standard SWC format and storing them accordingly. Additionally, the data is mapped to the same atlas space, and rich metadata is extracted to enhance the dataset’s utility. **C,** NeuroXiv has established the largest and most comprehensive dataset of neuron types, covering a wide range of brain regions including the TH, STR, Isocortex, HPF, and CB. A full list of abbreviations for all brain structures in this study is provided in **Methods**.

**Fig. 2.**
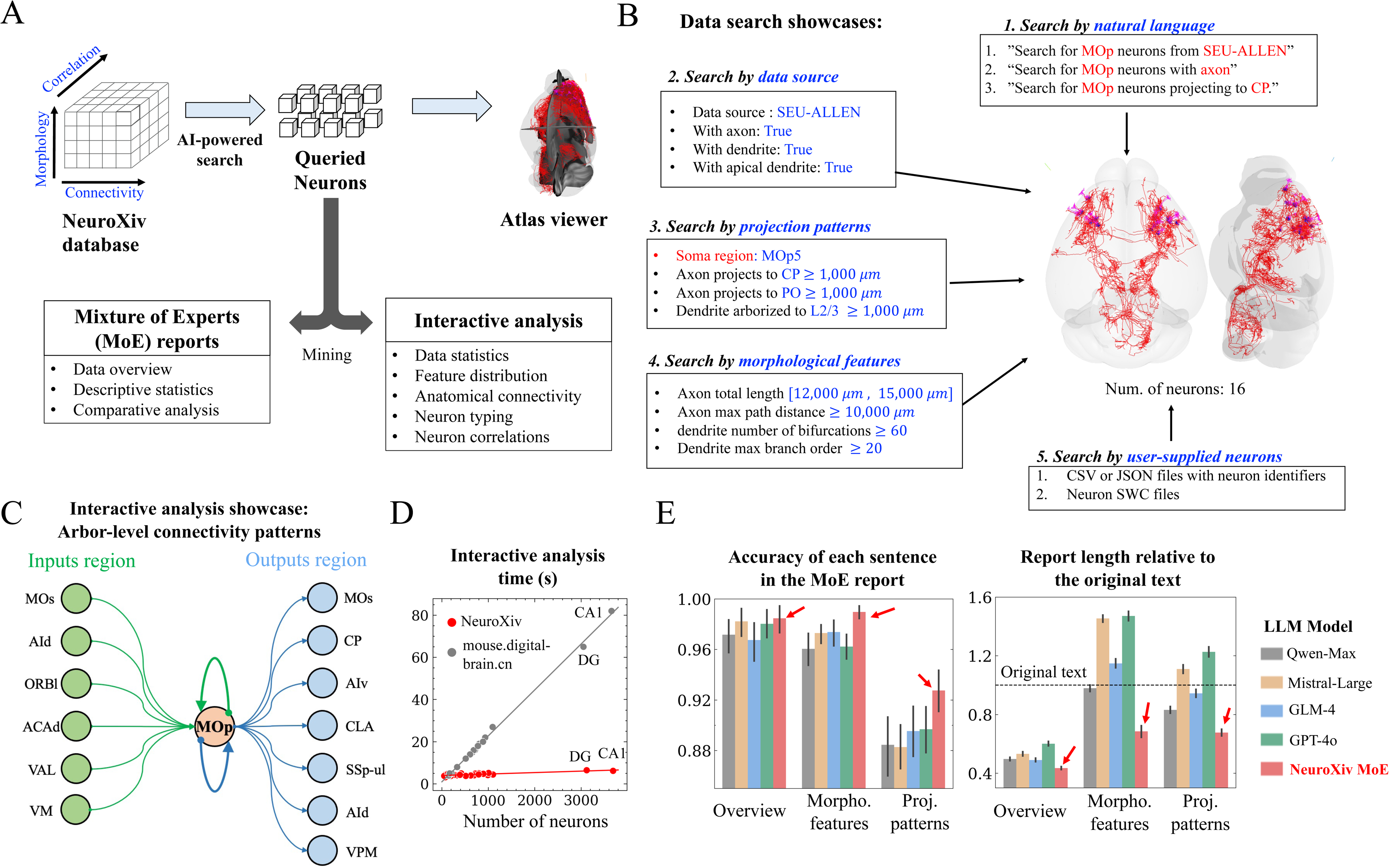
AI-powered engine for open analysis and mining of neuron data. **A**, the schematic diagram of data analysis and mining within NeuroXiv. The NeuroXiv database provides extensive morphology data, along with detailed morphological features, connectivity patterns, and inter-data correlations. The AI-powered search tool assists users in extracting relevant data of interest. Users can then interactively visualize the retrieved data on an atlas viewer and explore data features in depth. Additionally, NeuroXiv includes a Mixture of Experts (MoE) module that automatically generates reports to describe data characteristics and uncover patterns within the data. **B,** the showcases of data search in NeuroXiv demonstrate five common use cases. Users can search data using natural language queries, followed by searches based on the distribution of specific features provided by the user. Additionally, data can be retrieved using a user-supplied list of neurons, further enhancing the flexibility of data searches. **C,** the interactive analysis showcase in NeuroXiv presents an example where users can study arbor-level connectivity patterns. The input and output regions are determined by the spatial proximity of neurons within the database and the retrieved data, allowing for detailed exploration of neuronal connectivity. **D,** the comparison of interactive analysis time between the two platforms focuses on a shared cell type analysis scenario. The time measurement includes the entire process from data retrieval to the rendering of analysis charts. **E,** MoE reports were evaluated against four LLMs (Qwen-Max, Mistral-Large, GLM-4, and GPT-4o) for accuracy and text length across 100 random analysis cases. Our MoE showed higher accuracy and conveyed the same information using shorter text.

We initiate by establishing a server-side data processing pipeline capable of continuously aggregating publicly available datasets into our database (**Fig. 1B**). Interoperability is maintained through standardization of neuron morphology in the widely used SWC format (Mehta et al., 2023). Reusability is ensured by initially mapping neuron morphologies into the Common Coordinate Framework Version 3 (CCFv3) (Q. Wang et al., 2020), a widely recognized brain atlas for registering neuroanatomical data. We systematically document comprehensive metadata, encompassing basic information, morphological features, and anatomical arborization characteristics of neuron morphology (**Methods and Extended Data Table 2**). All morphology data and their corresponding metadata are accessible and downloadable through our web portal. Additionally, we have established an independent data server (download.neuroxiv.org) on AWS to expedite the download of entire standardized datasets.

We demonstrate the feasibility and scalability of the databasing method by consolidating data from diverse sources, including our SEU-ALLEN datasets (Peng et al., 2021; Y. Liu et al., 2023) and third-party examples (Winnubst et al., 2019; Gao et al., 2022, 2023; Qiu et al., 2024), culminating in the largest database of brain-wide neuron morphologies to date (**Fig. 1A, Methods** and Extended Data Fig. 1). The database features over 175,149 atlas-oriented reconstructed morphologies of individual neurons derived from more than 518 mouse brains. Each neuron reconstruction is characterized by its structural components (soma, axon, and dendrite), alongside metadata documented using a common standardized description method (**Supplementary Table 1**). The resource provides access to the most comprehensive atlas of cell types based on soma anatomical locations, encompassing 12 major gray matter divisions and including data from 292 out of 316 brain structures in the Allen Reference Atlas (ARA) ontology (**Fig. 1C**).

Our database offers several advantages for neuron morphology research. By mapping neuron morphologies from diverse sources onto a unified coordinate system, it enables rapid access to neuronal data without the need to switch between different sources. This data aggregation facilitates detailed, data-driven analyses of neuronal morphological characteristics and enhances the study of brain connectivity at the single-cell level (L. Liu et al., 2023). Specifically, we have identified a greater number of incoming neurons that extend their arbors into specific brain regions (Extended Data Fig. 3A) and have uncovered additional projection combinations of target regions formed by individual neurons (Extended Data Fig. 3B). Additionally, the database’s enhanced indexing system allows for improved retrieval of neurons based on spatial proximity, morphological similarity, or shared arborization patterns (Extended Data Fig. 2 and Extended Data Fig. 3C-D). This advanced indexing opens up new research opportunities, such as investigating whether spatially adjacent neurons consistently share similar morphological or projection characteristics (Extended Data Fig. 2A-B).

The AIPOM engine streamlines the knowledge development workflow on the established neuron morphometry database by integrating large language models (LLMs) (OpenAI et al., 2024; Touvron et al., 2023; Yang et al., 2024), which have rapidly advanced in recent years and demonstrated effectiveness in various domains (Bzdok et al., 2024; “Embedding AI in Biology,” 2024) due to their robust text comprehension capabilities. With AIPOM, users can flexibly define their data cohort of interest using natural language queries or a rule-based search panel. The LLM-based mining tool automatically transforms queried neuron data into comprehensive reports, including data overviews, descriptions of morphological features and connectivity patterns, and comparative analyses among cell types. Simultaneously, NeuroXiv provides an interactive analysis tool capable of generating a wide array of quantitative results for queried neurons, including the distribution of data attributes, morphological characteristics, and anatomical projection patterns through detailed visualizations (**Fig. 2A and Supplementary** Fig. 1). Additionally, NeuroXiv integrates an enhanced visualization tool for the interactive exploration of complex tree-like neuronal structures (**Methods, Extended Data** Fig. 8**, and Supplementary** Fig. 6-7).

We demonstrate the board applicability and flexibility of data search tool through several showcases of querying MOp neurons (**Fig. 2B**). We first illustrate that searches can be conducted using an LLM-based method, enabling users to query for specific neuron types, neurons with particular structures (such as axons or dendrites), or those exhibiting specific projection patterns through natural language inputs (**Supplementary** Fig. 2). We then show that precise searches can be performed by setting customized criteria based on neuron metadata (Extended Data Fig. 4A-C and **Supplementary** Fig. 3). NeuroXiv further provides a database interface enabling users to index specific neurons via an upload function, facilitating its use as a downstream exploration tool following user-defined neuron classification (**Supplementary** Fig. 4). Moreover, the search tool supports advanced capabilities such as similarity searches to identify neurons with comparable morphological features and arborization patterns (Extended Data Fig. 2A-B and **Supplementary** Fig. 8), as well as neighboring neuron queries to explore arbor-level connectivity within the brain (Extended Data Fig. 2C-D and **Supplementary** Fig. 9).

We highlight the interactive analysis capabilities of AIPOM through two studies. In the first study, users can identify which neuron types in the database provide input to a single neuron and determine the brain regions that receive projections from that neuron (**Fig. 2C**). For the second study, focusing on projection patterns, we use VPM neurons from our database as an example (Extended Data Fig. 9). These VPM neurons, sourced from two datasets, exhibit axonal arbors that extend across the CP into multiple cortical regions, including MOs, MOp, SSp, and SSs, encompassing various projection subtypes (Extended Data Fig. 9A). Additionally, our visualization tools allow users to observe the selectivity of different projection subtypes across cortical regions and layers (Extended Data Fig. 9B-C), as well as compare soma distribution and morphological features among these subtypes (Extended Data Fig. 9D-E). These findings align with prior knowledge of VPM neuron projection patterns (Peng et al., 2021; Y. Liu et al., 2023).

NeuroXiv also demonstrates high efficiency in online analyses (**Fig. 2D**). We perform benchmark tests comparing NeuroXiv and the Digital Brain platform by analyzing the same data categories and measuring the time to render results. Despite generating a broader range of analyses than the Digital Brain platform, NeuroXiv completes most tasks within 4-5 seconds and is largely unaffected by increases in data volume. In contrast, the Digital Brain platform exhibits sensitivity to data size, with response times increasing linearly. For example, generating results for the same number of CA1 neurons takes over 80 seconds on the Digital Brain platform, nearly 20 times slower than NeuroXiv.

To improve the integration of LLMs into AIPOM, we implement two key optimizations. First, to address the challenges of unpredictable outputs and occasional inaccuracies (Jin et al., 2024), we developed an advanced Mixture of Experts (MoE) framework for more reliable mining reports (**Fig. 2A and Extended Data** Fig. 5-7). This framework operates in three stages: first, a program generates standardized reports that capture all relevant data details; second, multiple LLM experts analyze and summarize these reports from a data scientist’s perspective; and third, a separate LLM reviews the outputs for accuracy and consistency, producing the final report. This multi-expert approach allows MoE to deliver comprehensive data overviews while effectively identifying morphological and projection differences. Our tests show that the MoE framework yields higher accuracy and more concise reports compared to those generated by a single LLM (**Fig. 2E and Methods**).

Second, to address the computational demands of server-side LLM deployment, we offer a client-side solution using a natural language processing (NLP) model and a supervised decision tree. This approach transforms natural language queries into actionable search operations within 2-3 seconds, achieving comparable accuracy with an 12.3-fold improvement in response time compared to LLM-based server-side searches (**Methods and Extended Data Table 3**).

In summary, NeuroXiv offers neuroscientists worldwide access to the largest and most comprehensive neuron morphometry resources. It aggregates publicly available neuron datasets from diverse sources, standardizes them into the widely used SWC format, and maps them into the CCFv3 atlas to enhance data reusability. Additionally, NeuroXiv integrates various tools, including search, visualization, and analysis, to facilitate rapid knowledge development. Leveraging advanced LLMs, the platform offers intuitive search functions and generates mining reports, thereby streamlining the extraction of valuable insights from neuron morphology data. To optimize LLM performance, AIPOM employs two approaches: the MoE framework for enhanced report accuracy and a client-side deployment for faster query responses. Together, the databasing method and AIPOM engine create an open, scalable, efficient, and flexible platform for ongoing neuron data reuse in the neuroscience community.

## Methods

### Nomenclature and abbreviations of brain regions

The 12 “major divisions” of gray matter in the Allen Reference Atlas (ARA) ontology: Isocortex, Olfactory areas (OLF), Hippocampal formation (HPF), Cortical subplate (CTXsp), Striatum (STR), Pallidum (PAL), Thalamus (TH), Hypothalamus (HY), Midbrain (MB), Pons (P), Medulla (MY), and Cerebellum (CB).

Isocortex: primary motor area (MOp), secondary motor area (MOs), primary somatosensory area (SSp), supplemental somatosensory area (SSs), gustatory area (GU), visceral area (VISC), dorsal auditory area (AUDd), primary auditory area (AUDp), posterior auditory area (AUDpo), ventral auditory area (AUDv), primary visual area (VISp), anterior cingulate area, dorsal part (ACAd), anterior cingulate area, ventral part (ACAv), prelimbic area (PL), infralimbic area (ILA), orbital area, lateral part (ORBl), orbital area, medial part (ORBm), orbital area, ventrolateral part (ORBvl), agranular insular area, dorsal part (AId), agranular insular area, posterior part (AIp), agranular insular area, ventral part (AIv), retrosplenial area, ventral part (RSPv), temporal association area (TEa).

### Olfactory areas (OLF): piriform area (PIR)

Hippocampal formation (HPF): hippocampal region (HIP), fields CA1, CA2, CA3, dentate gyrus (DG), entorhinal area, lateral part (ENTl), entorhinal area, medial part (ENTm), parasubiculum (PAR), postsubiculum (POST), presubiculum (PRE), subiculum (SUB), prosubiculum (ProS).

### Cortical subplate (CTXsp): claustrum (CLA)

Cerebral nuclei (CNU): striatum (STR), caudoputamen (CP), nucleus accumbens (ACB), globus pallidus, external segment (GPe), globus pallidus, internal segment (GPi). Thalamus (TH): ventral anterior-lateral complex (VAL), ventral medial nucleus (VM), ventral posterolateral nucleus (VPL), ventral posterolateral nucleus, parvicellular part (VPLpc), ventral posteromedial nucleus (VPM), ventral posteromedial nucleus, parvicellular part (VPMpc), medial geniculate complex, dorsal part (MGd), lateral geniculate complex, dorsal part (LGd), lateral posterior nucleus (LP), posterior complex (PO), anteromedial nucleus (AM), mediodorsal nucleus (MD), submedial nucleus (SMT), paraventricular nucleus (PVT), reticular nucleus (RT).

### Hypothalamus (HY): subthalamic nucleus (STN), zona incerta (ZI)

Midbrain (MB): substantia nigra, reticular part (SNr), midbrain reticular nucleus (MRN).

### NeuroXiv platform

The architecture of NeuroXiv (neuroxiv.org) is designed to support large-scale analysis of brain-wide neuron morphometry, utilizing a cohesive and highly integrated technology stack.

### Frontend

The frontend of NeuroXiv is developed using *Vue.js (v2.6.12)*, a progressive JavaScript framework known for its efficiency in building dynamic and responsive single-page applications. To create a user-friendly and visually engaging interface, *Element Plus (v2.7.3)*, a Vue 3-based component library, is employed. Additionally, *Three.js (v0.134.0)* is integrated into the frontend to handle the rendering of complex 3D visualizations, including brain regions and neuron reconstructions. This powerful WebGL-based library allows for detailed and interactive 3D models, providing users with an immersive experience in exploring neuroanatomical data.

### Backend

The backend is constructed with *Python (v3.9.12)* and *Flask (v3.0.0)*, a lightweight WSGI web application framework. Flask serves as the backbone of the server-side architecture, enabling seamless communication between the frontend and the database. *SQLite (v3.38.2)* is used as the database engine, offering a self-contained, serverless solution for efficient data storage and retrieval. This setup ensures that the platform remains agile and capable of handling the substantial datasets inherent to neuroinformatics research.

To manage web traffic and optimize performance, *Nginx (v1.24.0)* is deployed as a reverse proxy server. Nginx efficiently distributes incoming requests across backend processes, enhancing the platform’s ability to support a high volume of concurrent users while maintaining fast response times and secure connections.

NeuroXiv is hosted on Amazon Web Services (AWS) with 4 CPUs, 32 GB of memory, and 12.5 Gbps network bandwidth, leveraging AWS’s scalable and resilient cloud infrastructure to provide reliable access for users worldwide. This deployment strategy ensures that the platform remains highly available and capable of scaling in response to increasing user demand, thereby offering a stable and responsive environment for researchers. And we also noticed that the current performance of NeuroXiv on AWS is somewhat compromised and is expected to perform better on a server with the addition of more CPUs and acceleration through GPU devices.

### Datasets

Currently, the NeuroXiv database reports the integration of several brain-wide neuron morphology datasets shared by the community. Each dataset will be described in detail in the following sections. In the future, we plan to continuously add new mouse brain datasets and encourage users to contribute their own datasets to the NeuroXiv platform. Additionally, in upcoming updates, we plan to incorporate the BigNeuron Project (Manubens-Gil et al., 2023)— a community-contributed resource for benchmarking neuron morphology auto-tracing algorithms—into NeuroXiv, providing ongoing support for users worldwide.

### SEU-ALLEN full dataset

This dataset (Peng et al., 2021; Y. Liu et al., 2023) was initially generated using a semi-automatic annotation pipeline with 1,741 neurons and has been expanded to 1,876 neurons with improved quality (Li et al., 2023). Each neuron includes fully traced axonal and dendritic arbors, with 512 apical dendrites additionally annotated. The data mainly covers neurons in the VPM (389 neurons), CP (324 neurons), and lots of cortical regions.

### SEU-ALLEN local dataset

This dataset was produced using an automatic tracing method described in our previous work (Y. Liu et al., 2023). Initially, image volumes centered on the cell body (soma) were extracted from whole-brain image data, with a size greater than 200 µm in each dimension, sufficient to capture most of the neuron’s dendritic arbor. These image volumes were then processed using image enhancement algorithms (Guo et al., 2022) to improve image quality. Automatic reconstructions were generated and cross-validated using the APP2 (Peng et al., 2011; Xiao & Peng, 2013) and NeuTube (Feng et al., 2015) algorithms, followed by neurite fiber pruning to remove extraneous signals (Zhao et al., 2024). In NeuroXiv, we retained only the neurite segments within 100 µm of the soma to ensure consistency. This dataset contains 155,743 neurons, which are extensively distributed across various brain regions such as CP, MOB, OT, AON, and PIR.

### ION datasets

Currently, NeuroXiv integrates two datasets (Gao et al., 2022, 2023; Qiu et al., 2024) from ION: a prefrontal cortex dataset comprising 6,357 neurons (Xiaofei Wang. (2023). Single-neuron projectome of mouse prefrontal cortex (with dendrite). Brain Science Data Center, Chinese Academy of Sciences. https://cstr.cn/33145.11.BSDC.1689837400.1681922768243666945 and https://doi.org/10.12412/BSDC.1690164952.20001.), and a hippocampus dataset consisting of 10,100 neurons (Xiaofei Wang. (2024). Single-neuron datasets for mouse hippocampus. Brain Science Data Center, Chinese Academy of Sciences. https://cstr.cn/33145.11.BSDC.1667284058.1585980235450376194 and https://doi.org/10.12412/BSDC.1667278800.20001.). During data integration, 77 neurons with indeterminate soma locations were excluded, resulting in a final dataset comprising 16,380 fully reconstructed axons and 6,106 fully reconstructed dendrites. The neurons are primarily distributed across brain regions such as CA1 (3,657 neurons), DG-sg (2,618 neurons), SUB (934 neurons), and CA3 (887 neurons).

### MouseLight dataset

The MouseLight project (Winnubst et al., 2019) currently publishes data on 1,200 neurons available at MouseLight NeuronBrowser (http://ml-neuronbrowser.janelia.org). During data integration, 50 neurons with somas located in fiber tracts were excluded, resulting in 1,150 neurons from MouseLight being included in NeuroXiv. This dataset contains 1,150 fully reconstructed axons and 1,138 fully reconstructed dendrites. The neurons are distributed across various brain regions, such as MOs, SUB, PRE, VAL, DG-mo, and VPM, with some overlap with data from SEU-ALLEN and ION.

### Data aggregation

Data aggregation in NeuroXiv involves collecting datasets from our own datasets (SEU-ALLEN) and third-party sources like ION and MouseLight, and processing the data to convert it into a consolidated format.

### Data Format Conversion

SEU-ALLEN datasets have already been processed into the standardized SWC format (Mehta et al., 2023) and registered to the CCFv3 atlas. Therefore, our focus here is on processing the datasets from ION and MouseLight. The ION and MouseLight datasets had different format issues. We standardized the ION datasets into the SWC format, aligning the structure domain types, for example, soma (type label = 1), axon (type label = 2), basal dendrite (type label = 3), and apical dendrite (type label = 4). We also converted the neuron reconstruction data from the MouseLight dataset from JSON files into SWC files.

### Quality Control

We first performed a quality screening process to ensure data usability, filtering out non-compliant data. The specific steps included:

1. *Single Connected Tree:* Ensuring that all nodes have only one parent node, tracing back to a single root node (soma).
2. *Root Node (Soma) Labeling:* Verifying that there is exactly one node labeled as type=1 with parent=-1.
3. *Structure Domain Type Correctness:* Confirming that type attributes 1-4 are valid and that the type attribute remains consistent when tracing from terminal nodes back to the root.
4. *SWC Tree Structure Integrity:* Checking for the presence of loops and trifurcations in the SWC tree structure.

### Atlas Mapping

Using mBrainAligner (Li et al., 2022; Qu et al., 2022), we mapped all data points to the CCFv3 atlas. We then resampled the atlas-oriented reconstruction data, ensuring that the distance between parent and child nodes was set to 1 μm.

### Data Curation

All data points were renamed to follow a standardized format: “<resource_name>_<full/local>_<brainid>_<neuronid>_…_<atlas>”. For example: “SEU-ALLEN_full_17302_00001_CCFv3”.

### Metadata Extraction

We first extracted basic information such as the soma region for each neuron in the dataset. Then, using the atlas annotation template, we calculated the arborization strength for axons and dendrites across different brain regions based on neurite length. Finally, we extracted morphological features for axons and dendrites (Extended Data Table 2).

We further generated a list of morphology similar neurons for each neuron based on the distances between their morphological features. Additionally, we created a list of projection similar neurons by calculating the distance between point clouds formed by key axonal nodes (soma, bifurcation nodes, and terminal nodes). We also defined two types of neighboring neurons based on the overlap between neuron arbors: axon neighboring neurons and dendrite neighboring neurons. All metadata and proximity or similarity tables between data points have been imported into an SQL database for easy user access.

### Visualization

#### Brain Atlas Visualization

We use the Visualization Toolkit (VTK) to render brain atlases and neuron morphologies on the website. For brain atlas visualization, we start with annotation template files that record spatial coordinates for different brain regions, based on the ARA brain region table and brain templates. We apply the marching cubes algorithm to obtain the mesh contours for each brain region and generate corresponding mesh models. These models are then smoothed using the Laplacian smoothing algorithm. To optimize performance, we reduce the number of triangles in the mesh using the progressive mesh decimation algorithm while preserving the geometric information and other attributes. As a result, VTK files for 838 brain regions were generated for the CCFv3 atlas.

#### Neuron Morphology Visualization

This includes two components: a thumbnail view of each neuron morphology for the Neuron Browser and an interactive visualization in 3D atlas viewer for detailed neuron morphology. The thumbnails are designed to provide a quick overview of neuron morphology and assist users in locating neurons. To achieve this, we resample neuron morphologies with a step size of 100 µm. For generating 2D projection thumbnails, we first use Principal Component Analysis (PCA) to transform the neuron morphology coordinates, ensuring that the projection retains as much structural information as possible. For 3D visualization of neuron morphology, including axons, dendrites and arbors, we convert the neuron morphological structures into renderable line objects (OBJ files) for VTK. Additionally, soma visualization is implemented using Three.js, rendering a sphere with a radius of 50 µm.

### Mixture of Experts (MoE)

We have developed a Mixture of Experts (MoE) framework that leverages four large language models (LLMs), each containing trillions of parameters. This system is specifically designed to collaboratively mitigate errors and hallucinations that are commonly associated with LLM-generated content, thereby producing reliable, accurate, and coherent data analysis reports. The MoE framework operates in three distinct stages:

1 Descriptive reports generation: Initially, data retrieved from the database is programmatically organized into a standardized data description format. This ensures consistency and facilitates accurate analysis by the models.
2 LLM export reports: The organized data is then independently analyzed by three models— Qwen-Max-0428 (Yang, et al., 2024), Mistral-Large-2407 (AI, n.d.), and GLM-4-0520 (GLM et al., 2024). Each model is tasked with generating an analysis report based on the following prompts:

2.1 Prompt for Overview Objective: To provide a concise summary that enhances readability and clarity, with a focus on accurately representing significant numerical values. Methodology: The model is instructed to prioritize larger statistics while summarizing key findings in a coherent paragraph without bullet points. Data Input: “Original statistical data: {data}”
2.2 Prompt for Morphological Features Mining Objective: To analyze neuronal morphology data, particularly focusing on critical features such as ‘Total Length’ and ‘Number of Bifurcations.’ Methodology: The model generates a comparative summary that emphasizes the importance of these features, ensuring numerical accuracy throughout. Data Input: “Original neuronal morphology data: {data}”
2.3 Prompt for Projection Pattern Mining Objective: To analyze neuronal projection data, with a specific focus on axon and dendrite projections, and their implications for neuronal connectivity. Methodology: The model produces a summary that highlights the key points related to projection length and strength of connectivity, maintaining numerical precision and coherence. Data Input: “Original neuronal projection data: {data}”
3 Report confirmation: The GPT-4o-2024-05-13 model (OpenAI et al., 2024) serves as the final synthesis expert. This model evaluates the analysis reports generated by the three previous models against the original data and synthesizes them into a comprehensive, refined analysis report. The process follows a structured evaluation as outlined below:

3.1 Prompt for Overview Objective: To assess the precision of three summaries relative to the original statistical dataset insights. Methodology: The model ensures that numerical data in the summaries aligns with the source material. The most accurate summary is then refined into a new summary that enhances readability, brevity, and consistency. Data Input: “Original text: {origin_input}”
3.2 Prompt for Morphological Features Mining Objective: To meticulously assess the accuracy of three summaries in relation to an original text detailing neuronal morphology data. Methodology: The model compares numerical values, particularly those related to ‘Total Length’, ‘Number of Bifurcations’, ‘Max Path Distance’, and ‘Center Shift’, ensuring accuracy and consistency in the summaries. Data Input: “Original text: {origin_input}”
3.3 Prompt for Projection Pattern Mining Objective: To evaluate the precision of summaries concerning neuronal projection characteristics, particularly focusing on axon and dendrite projections as indicators of connectivity strength. Methodology: The model confirms numerical congruity and validates the logical consistency of comparisons in the summaries, generating a final, coherent summary. Data Input: “Original text: {origin_input}”

### MoE evaluation

The evaluation methodology is centered on assessing the accuracy and logical consistency of text summaries by comparing them against a source text. This process is implemented through a custom Python script that systematically evaluates key aspects of the summaries, particularly focusing on numerical data accuracy and logical consistency.

#### Data Accuracy Evaluation

The evaluation begins by extracting numerical data from both the source text and the generated summaries. A custom function utilizes natural language processing (NLP) tools, such as *spaCy*, to identify numbers within their contextual surroundings. These extracted numbers are then compared between the source text and the summaries to determine how accurately the numerical data has been represented.

A data accuracy score is calculated by examining the occurrence and contextual integrity of each number in the summaries relative to the source text. This score reflects the proportion of correctly matched numerical values, providing a quantitative measure of how faithfully the summaries represent the original data.

#### Logic Consistency Verification

Beyond numerical accuracy, the script also evaluates the logical consistency of the summaries. This involves verifying whether the statements in the summaries logically follow from the information provided in the source text.

The script employs a large language model (LLM) to perform this verification. It generates a prompt that includes both the source text and the summary statement in question, asking the model to determine whether the summary statement can be logically and numerically inferred from the source. The output from the LLM is then parsed to decide whether the summary is logically consistent. The logic accuracy score is derived by calculating the percentage of summary sentences that were deemed logically valid.

#### Comprehensive Evaluation

The script integrates the results from both the data accuracy and logic consistency assessments to provide a comprehensive evaluation of the summaries. By quantifying the alignment of numerical data and logical coherence, the evaluation method offers a robust approach to determining the quality and reliability of text summaries in capturing the essence of the source material.

#### Documentation and Reporting

The results of the evaluation process, including both data accuracy and logic consistency scores, are meticulously recorded. This documentation includes relevant metadata, such as the models used and the specific instances evaluated, ensuring that the evaluation process is both transparent and reproducible for further analysis and refinement.

#### AI-powered natural language search

Our framework integrates multiple components to achieve accurate and context-aware natural language understanding and data retrieval.

1. Entity Recognition and Intent Classification: The core of our Natural Language Processing (NLP) framework is built on a combination of machine learning models and rule-based systems. A supervised decision tree classifier, trained on a specialized dataset of neuroscience-related queries, is used to recognize key entities such as neuron types, brain regions, and projection relationships. The classifier works alongside rule-based components that handle domain-specific terminology variations, ensuring a robust response to user queries.
2. Semantic Parsing and Contextual Understanding: The framework employs semantic parsing techniques to accurately extract and interpret user intent from natural language input. It detects complex phrases related to neuroscience, such as neuron classifications and brain region relationships. Using contextual analysis, the system discerns detailed query intents (e.g., “projection from region X to region Y”), allowing precise and relevant data to be retrieved.
3. Dynamic Mapping and Knowledge Integration: The framework integrates domain-specific structured schemas to map both full terminologies and their corresponding abbreviations into a standardized format compatible with database queries. This dynamic mapping ensures consistency and accuracy by aligning user input with the system’s structured knowledge base. This capability enhances the system’s flexibility and robustness in providing comprehensive and relevant responses.
4. Multi-Stage Query Processing Pipeline: The NLP module operates through a multi-stage query processing pipeline, encompassing tokenization, entity extraction, context recognition, and result formulation. Each stage is designed to maximize the understanding of user input and generate accurate database queries, providing users with precise and comprehensive results.

#### Front-End Deployment and Benefits

The NLP framework is deployed on the front end using Vue.js, which brings two significant advantages:

1. Protecting User Privacy: By processing queries directly on the client side, the framework ensures that user inputs remain private and are not exposed to external servers. This approach is particularly beneficial in sensitive research settings where data privacy is paramount.
2. Improved Query Speed and Responsiveness: Client-side processing significantly reduces latency by eliminating unnecessary server round trips. This results in faster response times and a more interactive user experience, enabling researchers to explore neuroscience data efficiently.

#### Implementation and Model Training

The implementation leverages JavaScript-based libraries combined with tailored AI algorithms optimized for the neuroscience domain. The decision tree model is trained on a diverse set of domain-specific queries to ensure robust performance and generalization.

## Data availability

Atlas-mapped neuronal morphology data and discovery results—including metadata, figures, and mining text—are available for direct download via the web portal. Additionally, we have set up an AWS server (https://download.neuroxiv.org) to facilitate easy access to standardized datasets. VTK files for various brain regions in CCFv3 are also accessible through our GitHub repository (https://github.com/SEU-ALLEN-codebase/NeuroXiv/VTK). All the materials available from NeuroXiv should only be used exclusively for academic purposes and must adhere to the CC-BY NC license (https://creativecommons.org/licenses/by-nc/4.0/legalcode).

## Code availability

The source code of the NeuroXiv project can be obtained from our GitHub repository (https://github.com/SEU-ALLEN-codebase/NeuroXiv), where detailed deployment instructions are provided. User manuals and video-tutorials are available at our website (https://neuroxiv.org). Data processing utilizes the plugin system provided by the Vaa3D platform (version 4.001, available at https://github.com/Vaa3D). The source code for quality control procedures can be found at https://github.com/Vaa3D/vaa3d_tools/tree/master/hackathon/shengdian/NeuroMorphoLib. The source code for mBrainAlinger is accessible at https://github.com/Vaa3D/vaa3d_tools/tree/master/hackathon/mBrainAligner. Atlas mapping can also be conducted via the mBrainAlinger web portal (http://mbrainaligner.ahu.edu.cn).

## Supplementary Data

The Supplementary Notes (*NeuroXiv_supplementary_notes_and_figures.pdf*), Supplementary Tables (*Supplementary_Table1.csv*) and Supplementary Figures (*NeuroXiv_supplementary_notes_and_figures.pdf*) can be found along with the submission files of this manuscript.

## Acknowledgements

This work was mainly supported by several initiatives of neuroscience and a New Cornerstone grant awarded to H.P.. The Southeast University team was also supported by a STI2030-Major Projects Grant No. 2022ZD0205200/2022ZD0205204 awarded to Lijuan Liu. We thank Brain Science Data Center, Chinese Academy of Sciences (https://braindatacenter.cn/) for the open sharing of ION datasets. We thank MouseLight project (http://ml-neuronbrowser.janelia.org) for the open sharing of its dataset. We extend our gratitude to Lijuan Liu for her contributions to the project design and discussions surrounding the first-generation platform, and to Xuan Zhao for early involvement in the project’s development. We also thank Sujun Zhao for assisting with the use of the auto-arbor tool to generate arbor data for all axon-containing neurons, and Xiaoxuan Jiang for testing NeuroXiv’s functionalities and drafting the NeuroXiv user manual.

## Author contributions

H.P. conceptualized and managed this study, and invented AIPOM. Z.Y., H.C., and J.Y. designed the initial version of the NeuroXiv (neuroxiv.net) platform, hosted originally on Tencent Cloud server. Z.Y. was responsible for backend development of the first-generation platform, while H.C. handled frontend coding. S.J. and L.W. co-developed the new version of the NeuroXiv (neuroxiv.org) platform and migrated the servers to the AWS cloud platform. L.W. undertook the majority of website development tasks, including project deployment, backend, and frontend development. S.J. managed the data aggregation tasks, including data standardization processing, metadata generation, and the production of required atlases and reconstruction files (obj files) for the website. H.P. and S.J. wrote the manuscript with assistance of all authors, who reviewed and revised the manuscript.

## Extended Data Figures

**Extended Data Fig. 1.**
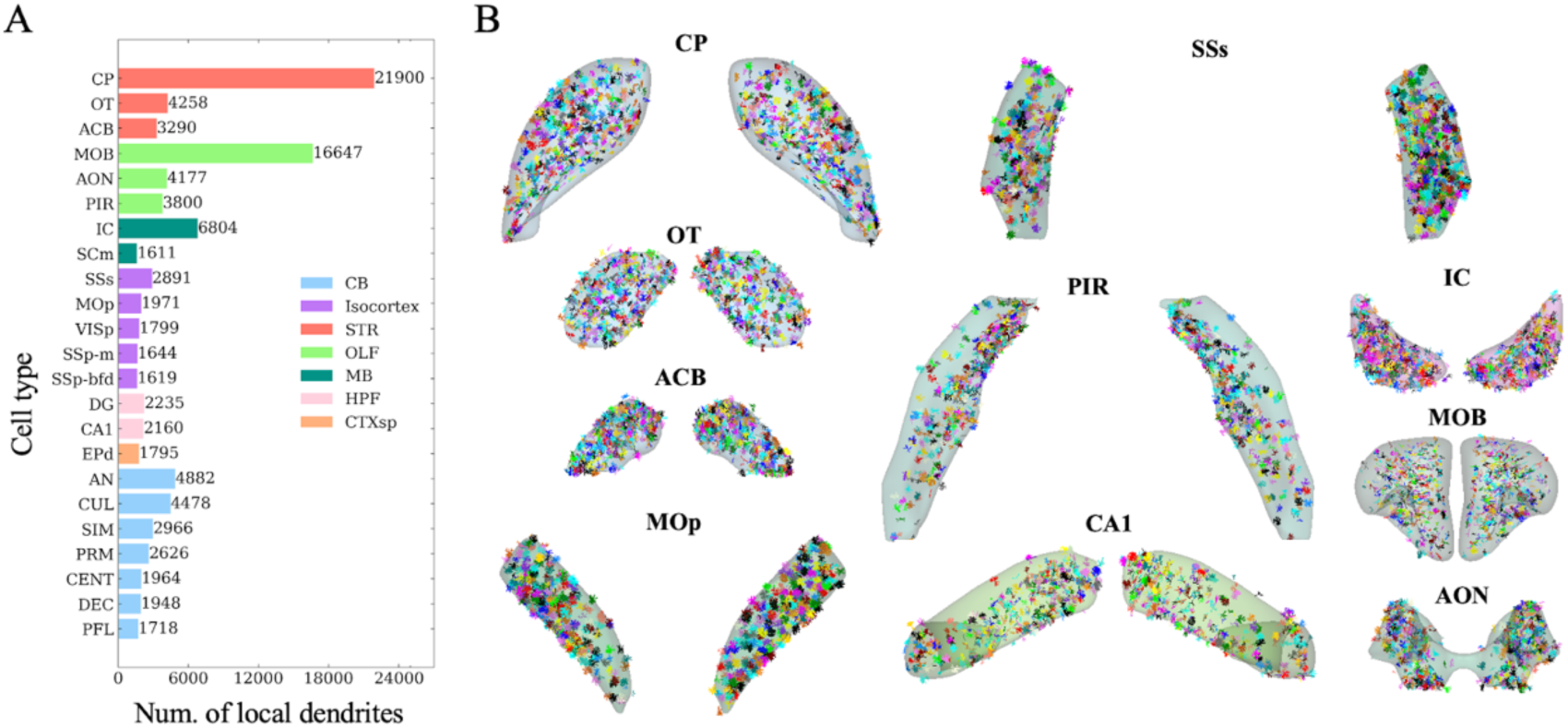
Local dendrites in the NeuroXiv database. A, the statistics of the number of local dendrites in different brain regions within the CCFv3 atlas. B, Visualization of local dendritic data across various brain regions. Local dendrites are rendered in distinct colors to enhance differentiation.

**Extended Data Fig. 2.**
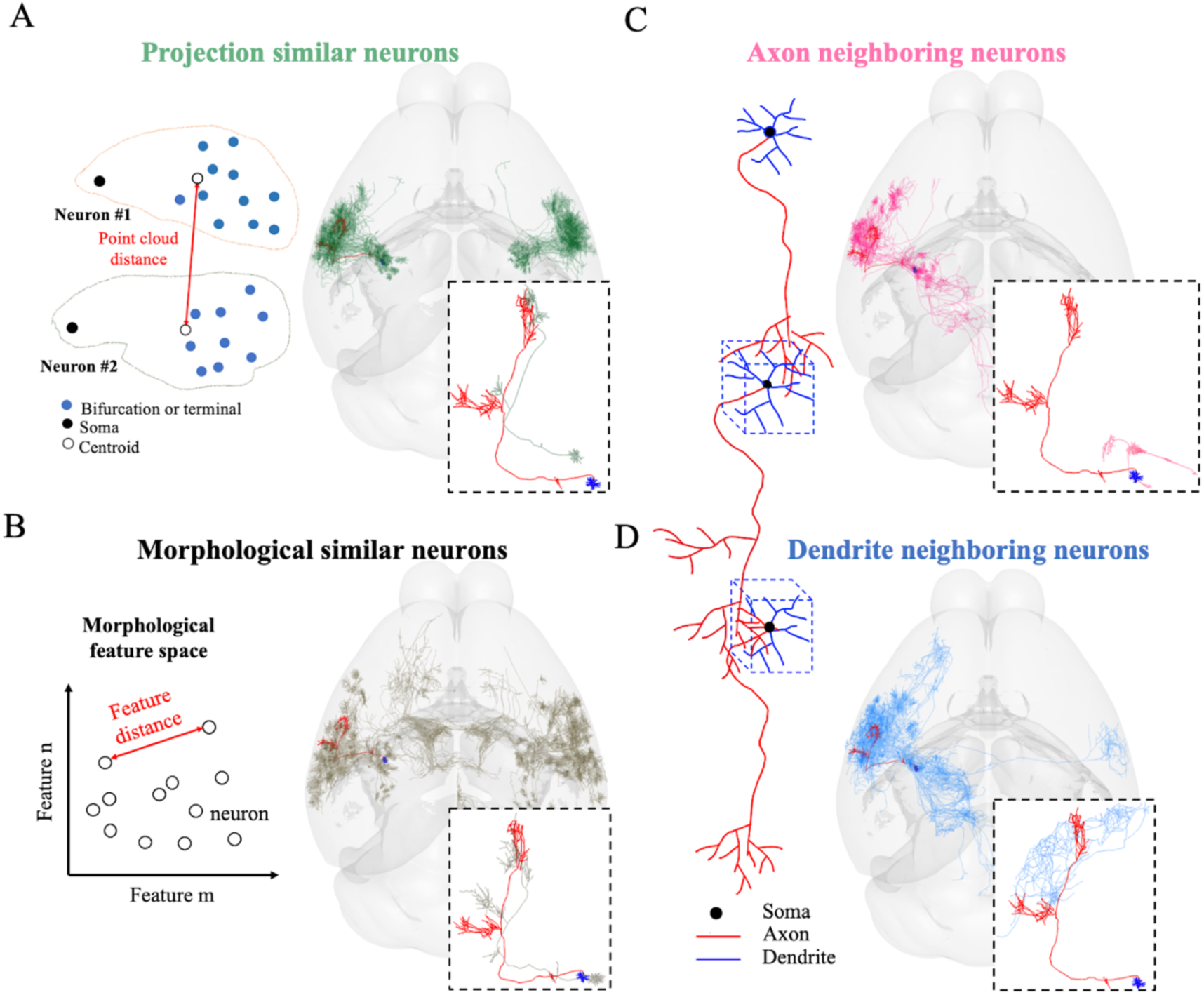
Illustrative diagrams depicting the definition of correlated neuron data on the NeuroXiv platform. A, projection similarity neurons are defined by the distance measured between key axonal nodes of the neurons, including the soma, bifurcation points, and terminal points. B, morphological similarity neurons are defined by calculating the distance between neuron pairs in morphological feature space. In the database, we rank similarity based on the distances, with closer distances indicating greater similarity. C and D, Axon neighboring neurons are those where the axonal arbor of neurons in the database spatially overlaps with the dendritic arbor of the subject neuron. Conversely, dendrite neighboring neurons are those where the dendritic arbor of neurons in the database spatially overlaps with the axonal arbor of the subject neuron. In the database, we rank neighboring neurons based on the length of the overlapping arbor regions, with greater lengths indicating higher proximity.

**Extended Data Fig. 3.**
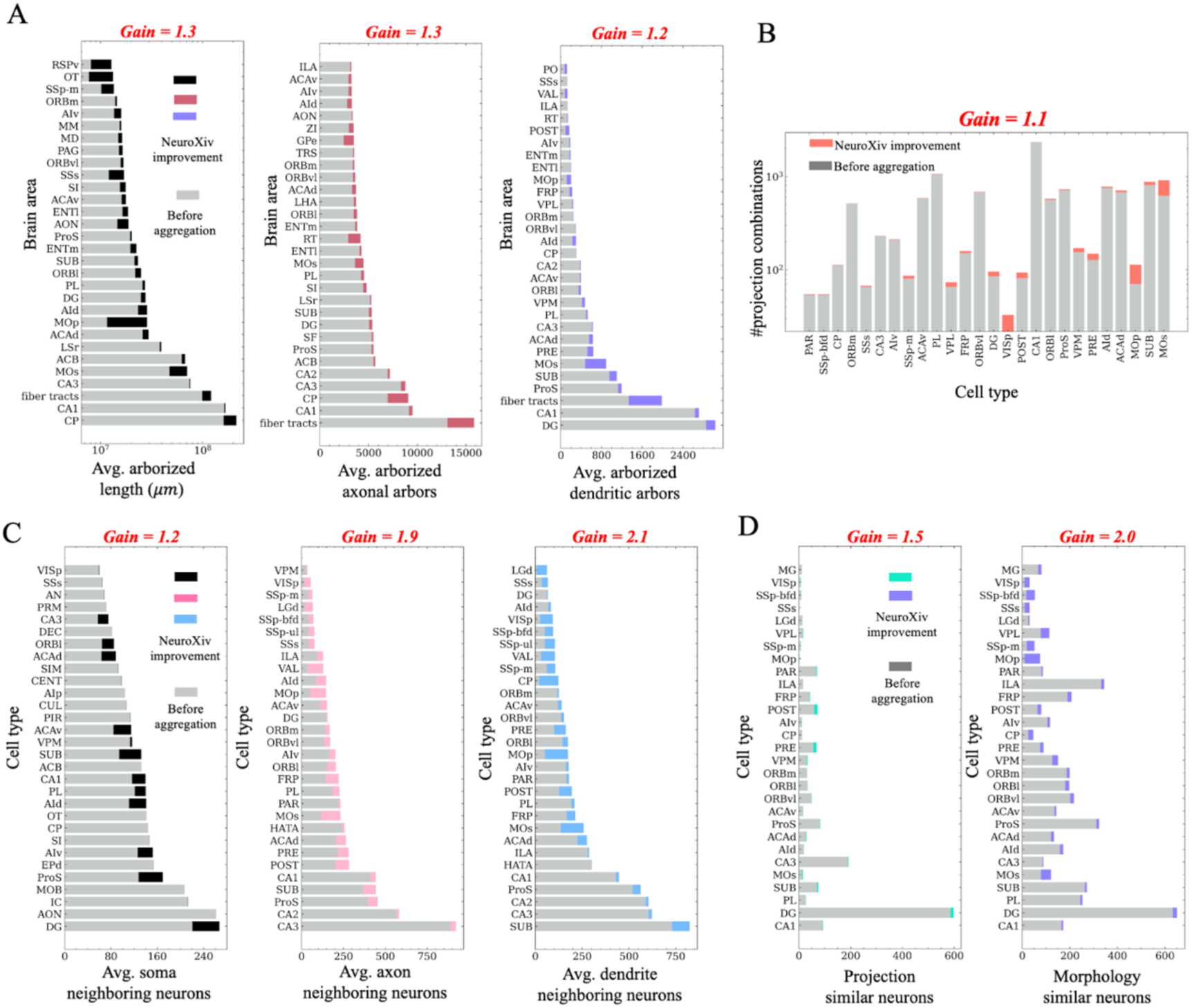
Gains in improvements resulting from data aggregation. **A**, we aggregated three data sources to formulate the NeuroXiv database. Compared to the information obtainable from a single data source, this approach allows us to capture more neurite length across various regions. The gray bar represents the maximum neurite length that can be obtained from one data source, while the other colored bars represent the improvements gained through aggregation. **B**, the NeuroXiv database offers a greater number of projection combinations, where neurons extend into brain regions with lengths exceeding 1000 μm. **C** and **D**, we can identify more neighboring neurons, including soma neighboring, axon neighboring, and dendrite neighboring neurons. Additionally, we can find more similar neurons, including those that are projection similar and morphology similar. Definitions of neighboring and similar neurons can be found in **Extended Data Fig. 2**.

**Extended Data Fig. 4.**
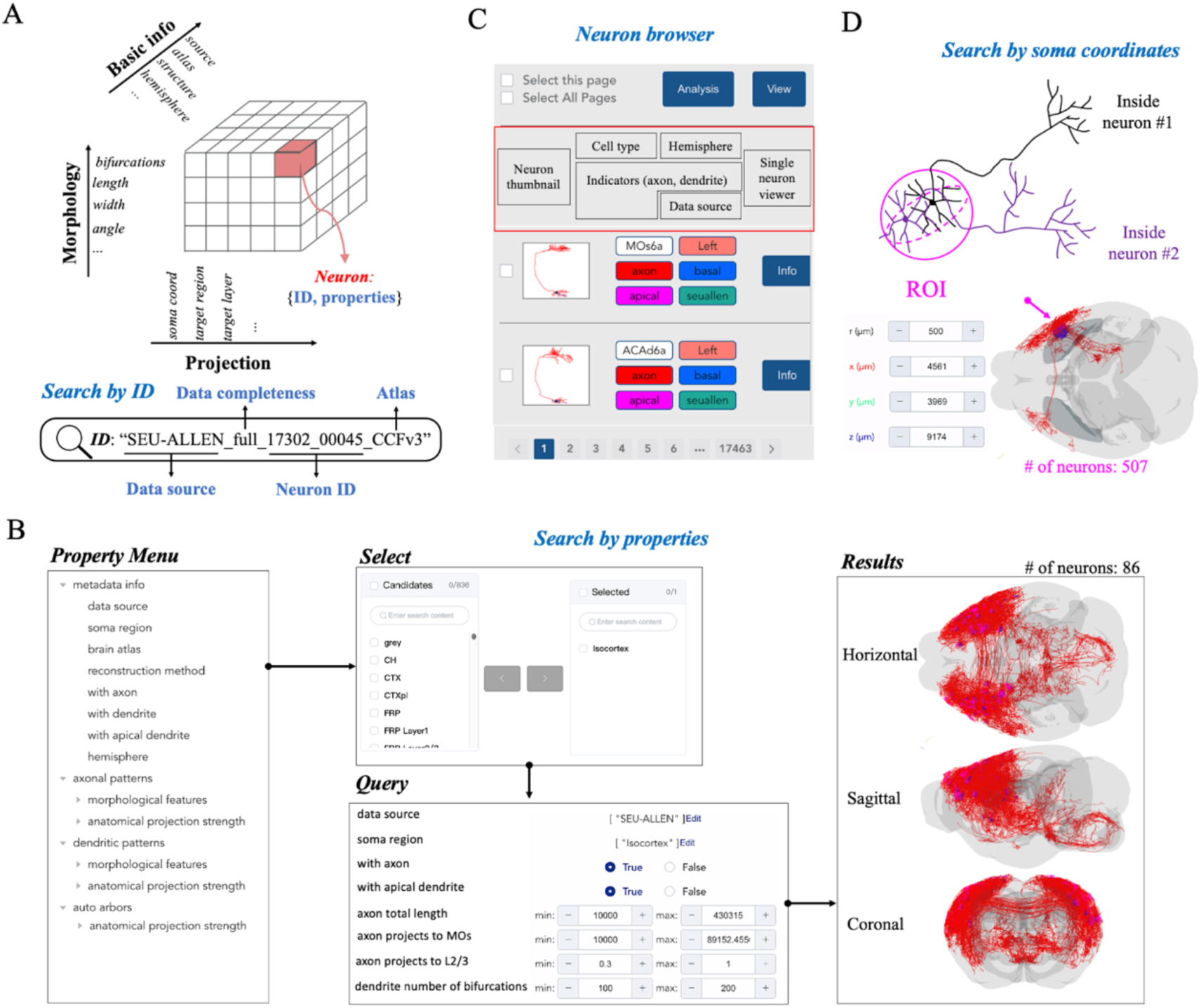
An illustrative diagram of data retrieval and filtering on the NeuroXiv platform. A, each neuron reconstruction in the NeuroXiv database is assigned a unique ID and includes three types of metadata: basic information, morphological features, and projection characteristics. B, users can customize their search strategies using either the ID or the metadata. C, to facilitate efficient data filtering, we have implemented a neuron browser on the web portal. This tool displays the morphology of each neuron (neuron thumbnail) and key information, and includes an entry point for navigating to detailed data pages. D, users can define regions of interest (ROI) and then retrieve neurons with soma located within these ROI (Supplementary Fig. 5).

**Extended Data Fig. 5.**
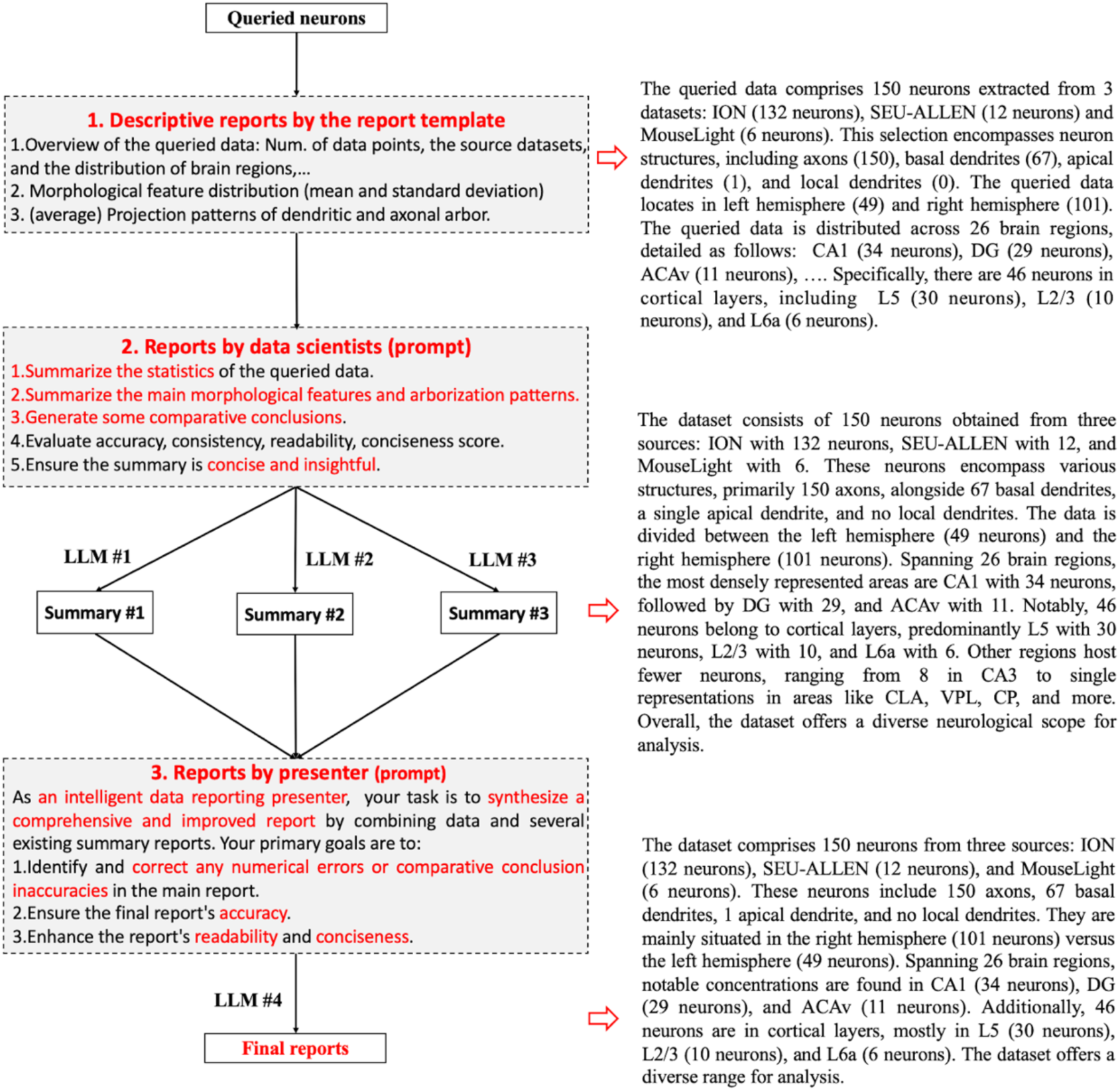
A schematic illustration of the mixture of experts (MoE) system. The MoE-based report generation process can be divided into three distinct stages. 1) Descriptive reports generation: A program generates reports in a fixed format, capturing all relevant details of the retrieved data. 2) LLM Expert reports: Multiple LLM Experts analyze and summarize the descriptive reports from a data scientist’s perspective. Although three experts are shown, the process can involve one or more. 3) Report confirmation: A different LLM Expert evaluates the previous reports for accuracy, readability, and coherence, and refines the final report accordingly. An actual case is shown on the far right, with red arrows pointing to the reports generated at each stage.

**Extended Data Fig. 6.**
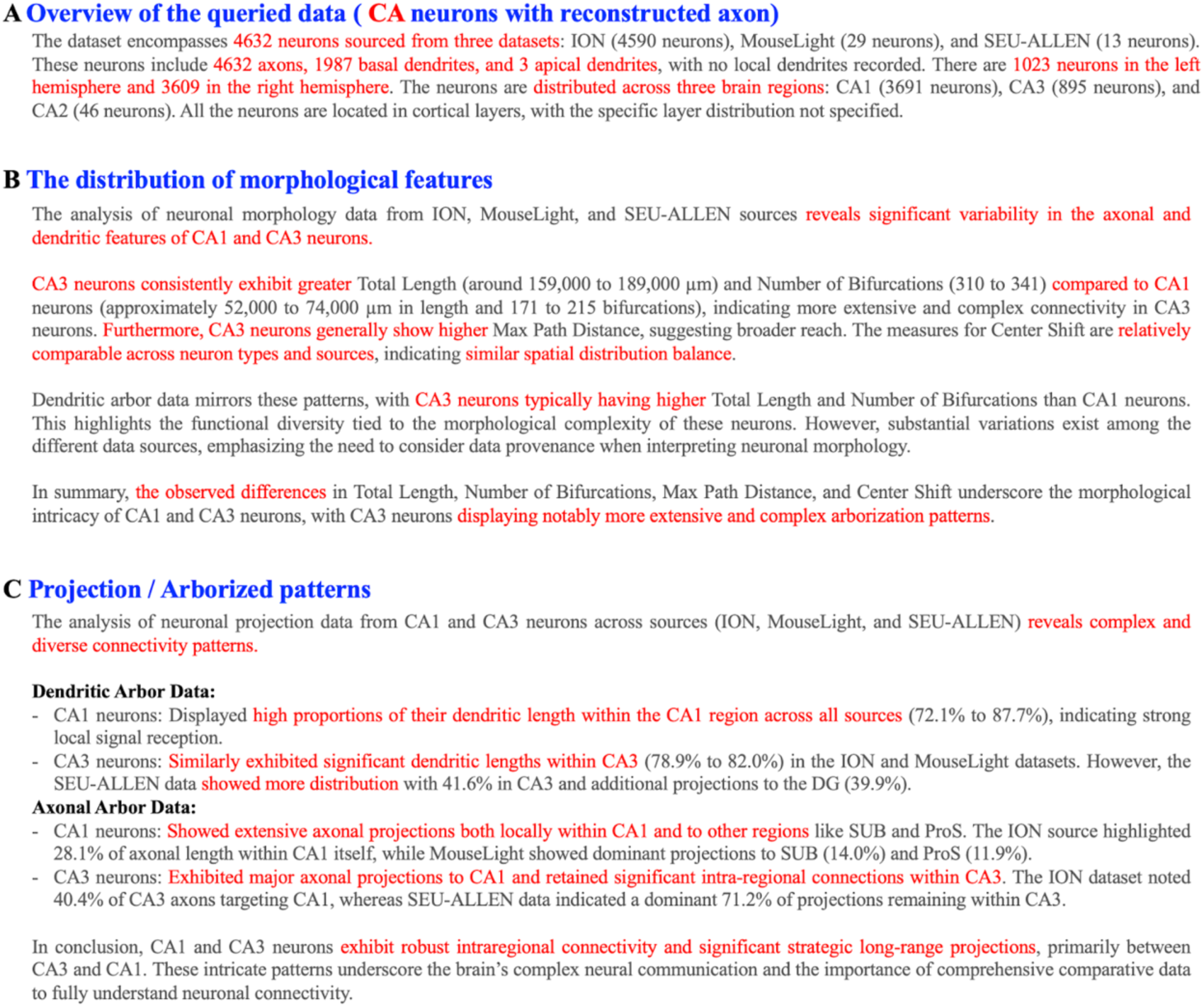
MoE report of the queried CA neurons. The report is structured into three sections: A) an overview of the queried data, B) the distribution of morphological features within the queried data, and C) the projection patterns observed in the queried data. Key statistical points and comparative analyses are highlighted in red throughout the report.

**Extended Data Fig. 7.**
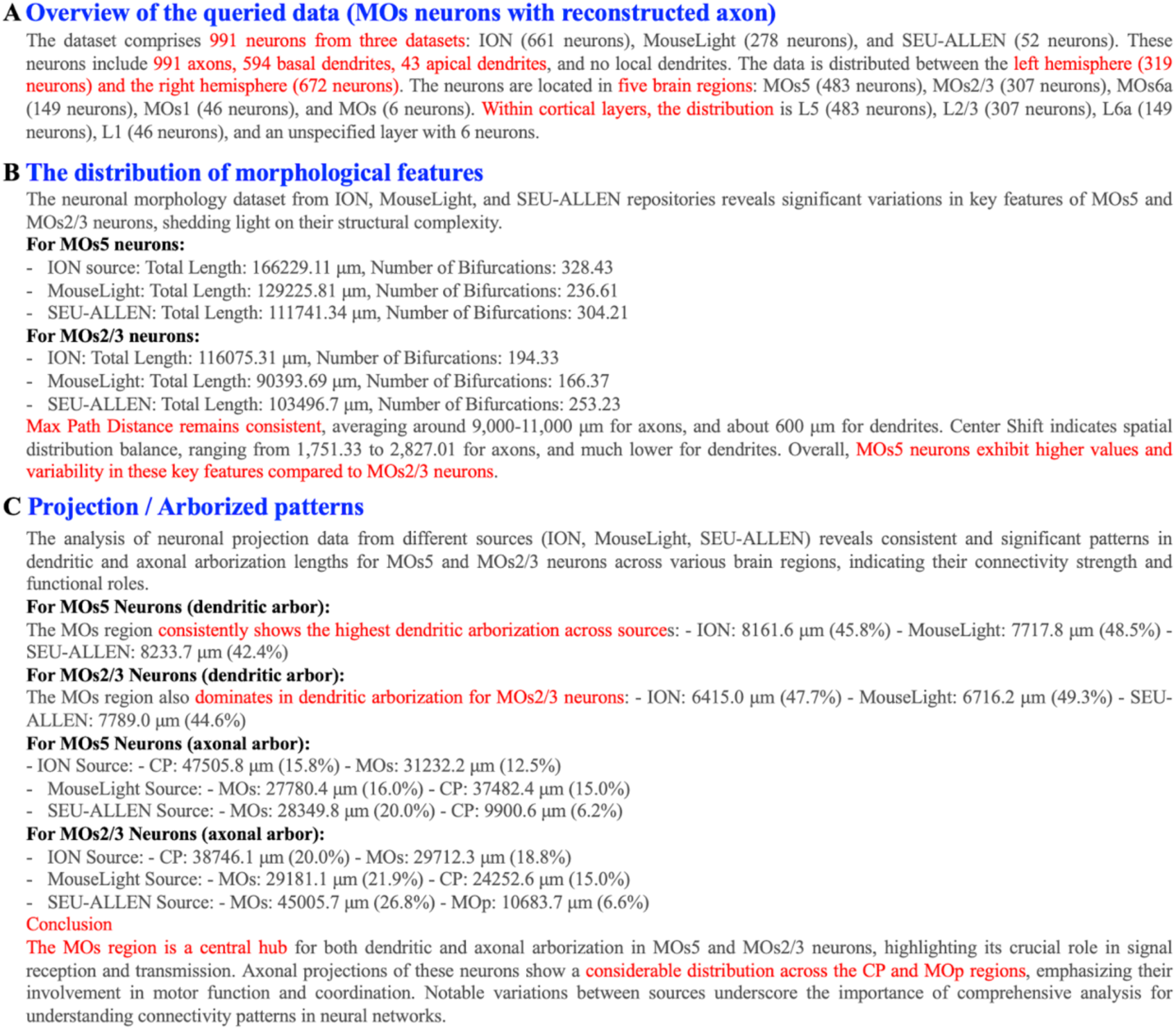
MoE report of the queried MOs neurons. The report is structured into three sections: A) an overview of the queried data, B) the distribution of morphological features within the queried data, and C) the projection patterns observed in the queried data. Key statistical points and comparative analyses are highlighted in red throughout the report.

**Extended Data Fig. 8.**
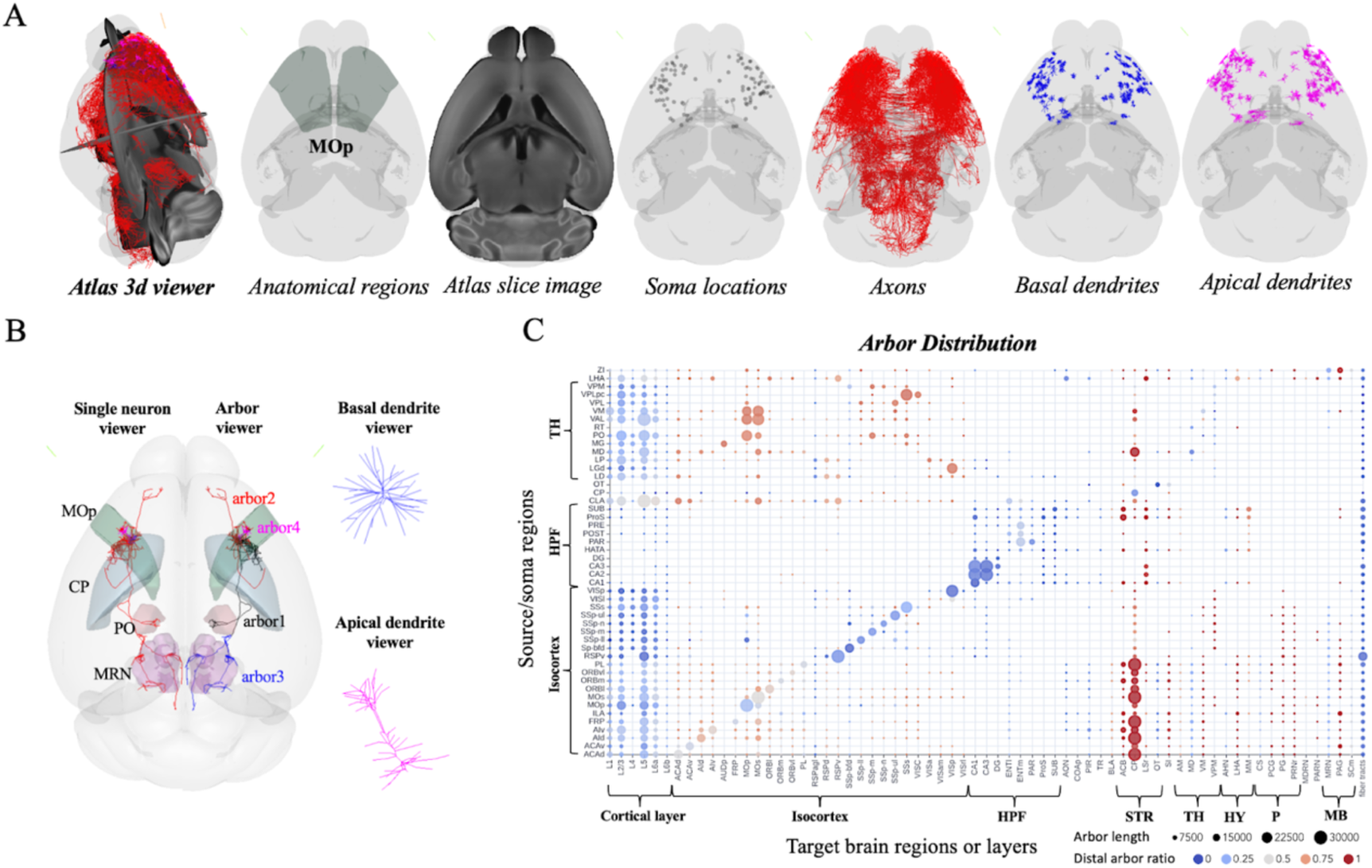
Interactive visualization diagram of NeuroXiv. **A**, we have implemented a 3D viewer on the web platform, allowing users to visualize the atlas and neurons or neuronal structures of interest. **B**, visualization of individual neuron data, featuring an embedded arbor viewer with zoom-in views of basal and apical dendrites. **C**, arbor distribution of major cell types across various brain regions. We also visualize arbor distribution across different cortical layers.

**Extended Data Fig. 9.**
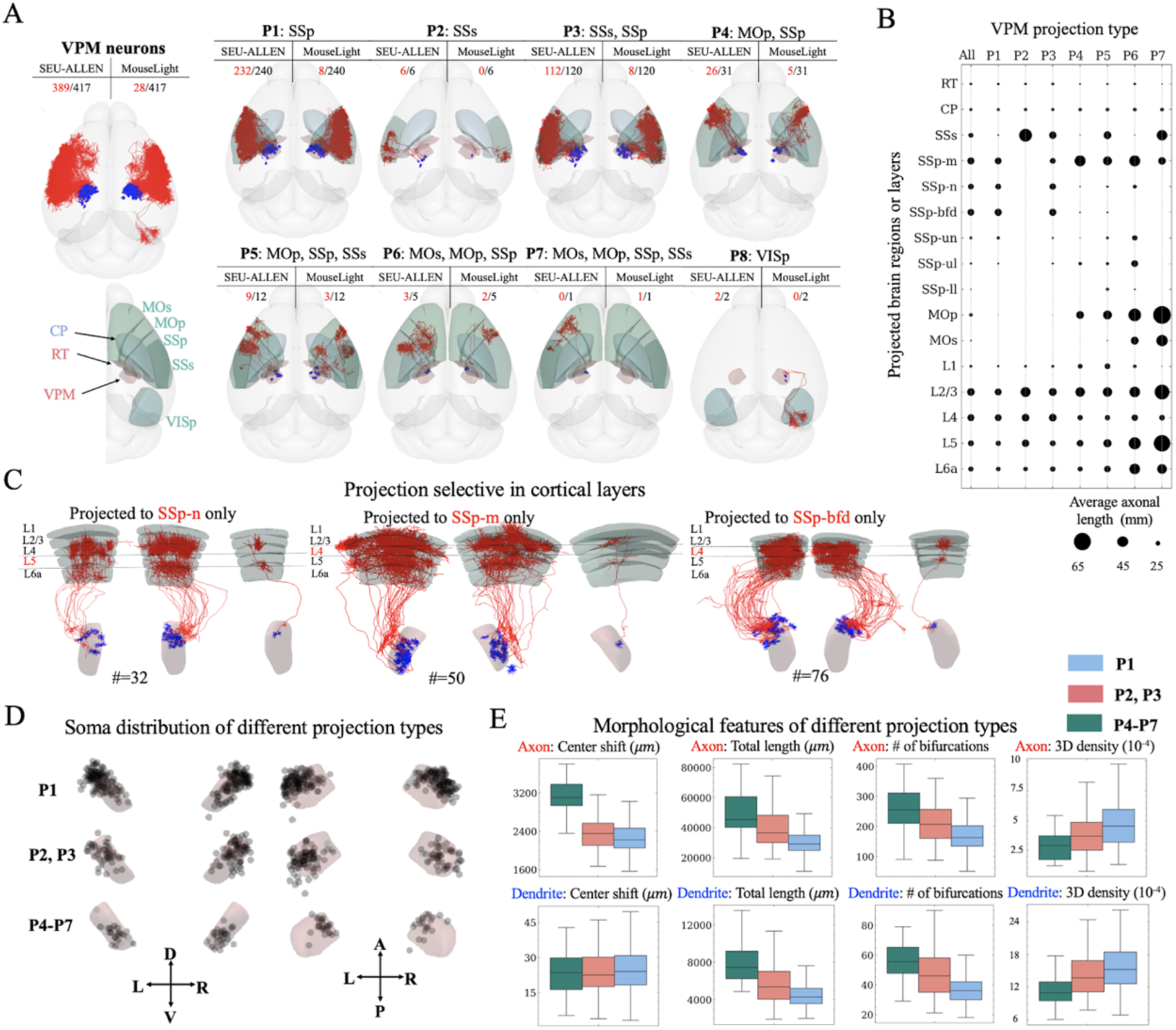
A case study of neuron projection research using NeuroXiv. **A**, this study includes 417 VPM neurons from two sources. Overall, VPM neurons project through the CP brain region, bifurcating at the boundary of the CP region to target cortical brain areas. Based on combinations of target brain regions, eight subtypes of projections are identified. Notably, neurons projecting to VISp (subtype P8) exhibit entirely unique projection patterns, which were excluded from the downstream analysis due to their distinctiveness. **B**, differences in projection strength among various projection types in target brain regions and cortical layers are visualized, with projection strength determined by axonal length. **C**, VPM neurons exhibit projection selectivity across cortical layers. For instance, neurons projecting solely to the SSp-n region form clusters while skipping L5, whereas neurons projecting solely to the SSp-m or SSp-bfd regions form clusters while skipping L4. **D** and **E**, differences in soma distribution and morphological characteristics among various VPM projection subtypes are analyzed.

## Extended Data Tables

**Extended Data Table 1.**
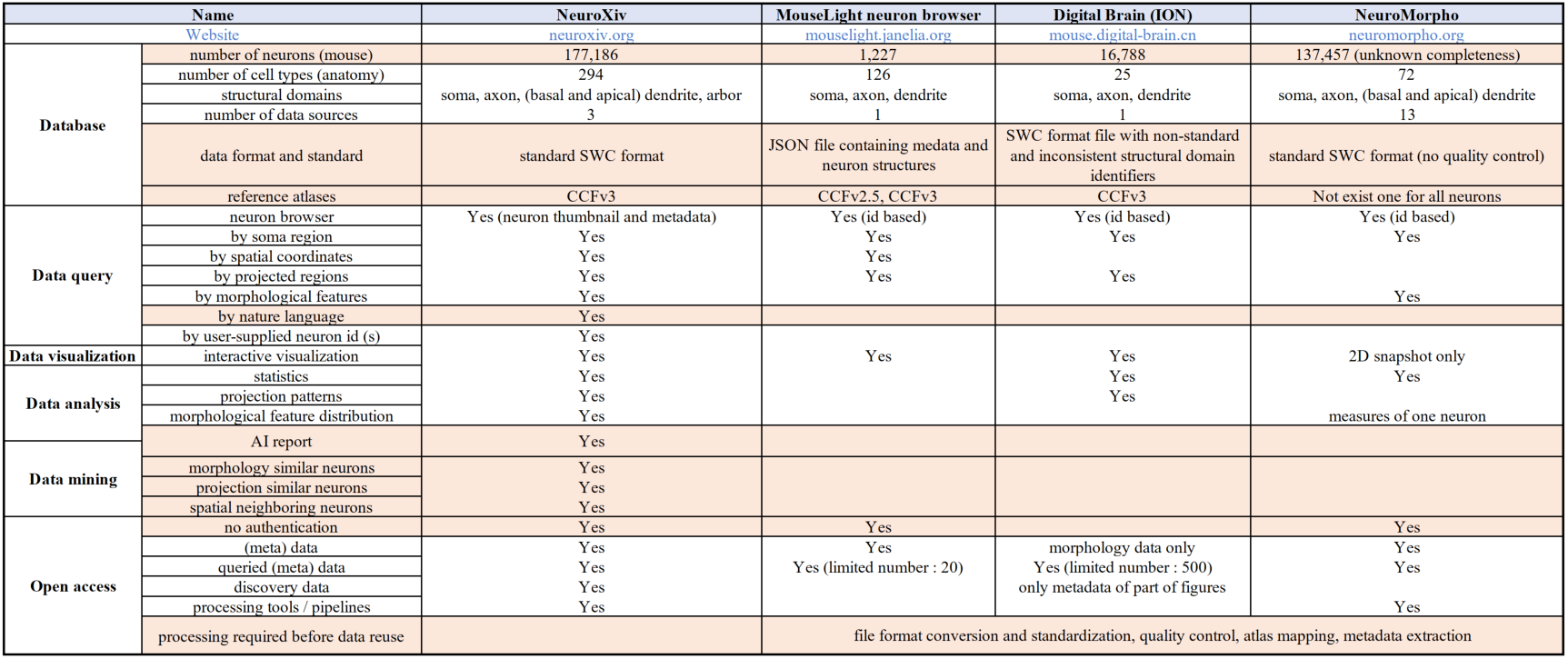
Comparison of alternative neuron morphology web platforms (*Extended_Data_Table1.pdf*).

**Extended Data Table 2** The metadata of neurons and their descriptions in the NeuroXiv database (*Extended_Data_Table2.pdf*). This table can be found along with the submission files of this manuscript.

**Extended Data Table 3.**
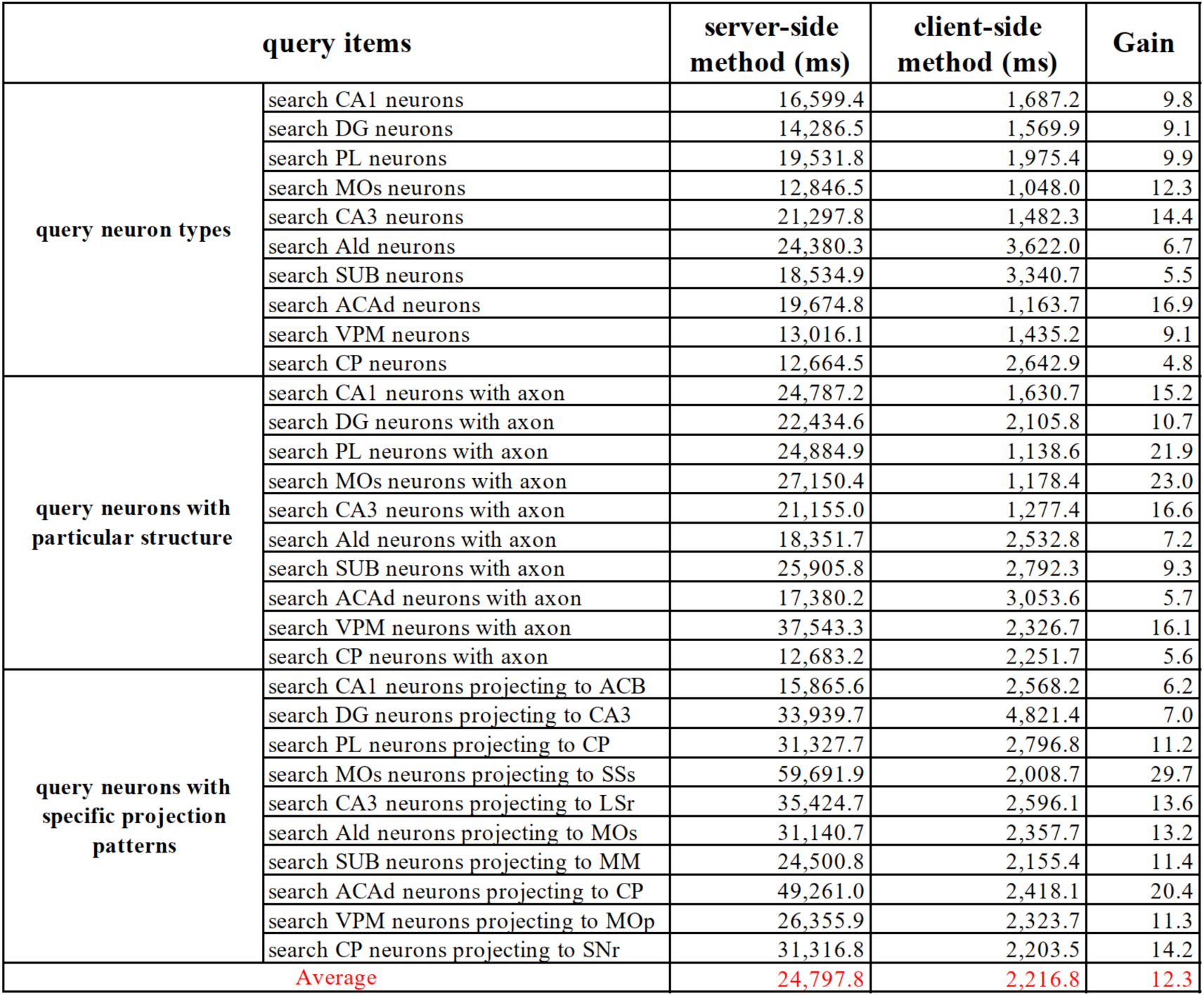
Comparison of the performance of two types of natural language query methods in AIPOM. (*Extended_Data_Table3.pdf*). To achieve this, we defined three common retrieval scenarios: querying neuron types, querying neurons with particular structures, and querying neurons with specific projection patterns, each comprising 10 test cases. To ensure fair testing, both methods used the same computer configuration, eliminating disparities due to varying computational power. The server-side method involved setting up a local NeuroXiv server on the test computer, while the client-side method accessed the NeuroXiv server hosted on AWS directly. As a result, compared to the server-side approach, the client-side method demonstrated an average response time improvement of 12.3.

