## Supplementary Notes for "Beyond Static Brain Atlases: AI-Powered Open Databasing and Dynamic Mining of Brain-Wide Neuron Morphometry"

### Supplementary Figures

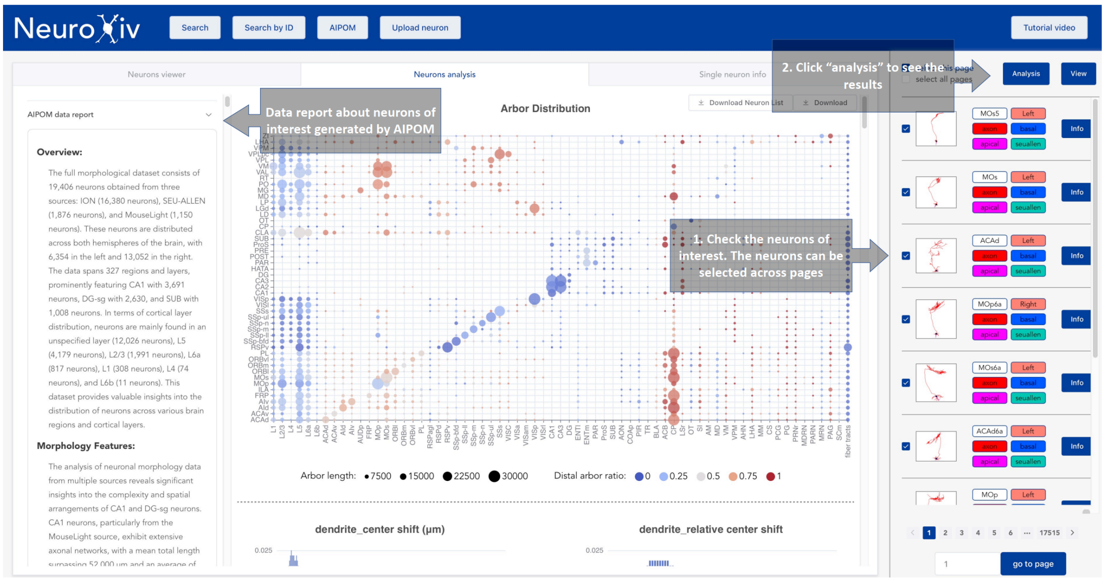


**Supplementary Fig. 1 | A screenshot of the neuron analysis panel.** *Users can select neurons of interest in the neuron browser panel for analysis, and obtain arbor distribution, feature distribution, and MoE reports directly from this panel.*

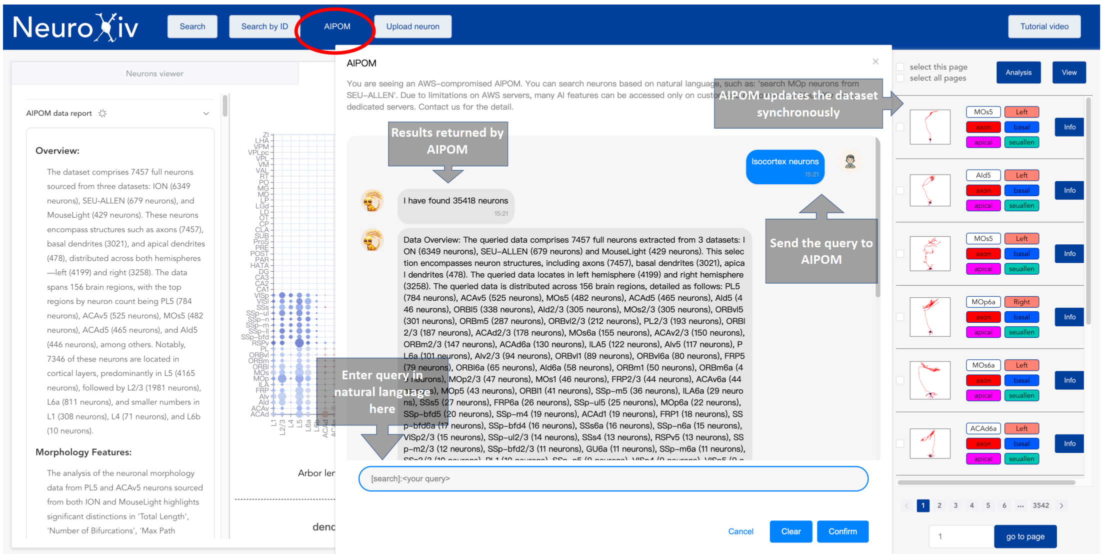


**Supplementary Fig. 2 | A screenshot demonstrating how to use natural language to search for neurons.** *A text box or input field allows users to enter their search queries. Upon entering the query, the interface automatically recognizes the intent to search for neurons. Once the intent is detected, the Neuron browser panel is automatically updated to display neurons that match the specified search criteria.*

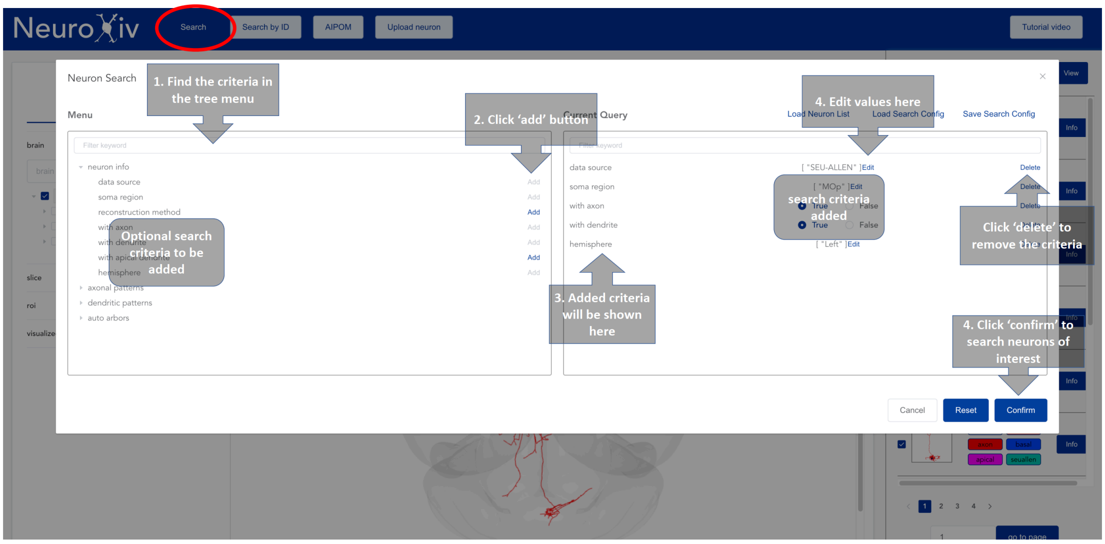


**Supplementary Fig. 3 | A screenshot demonstrating how to customize specific search criteria to search for neurons.**

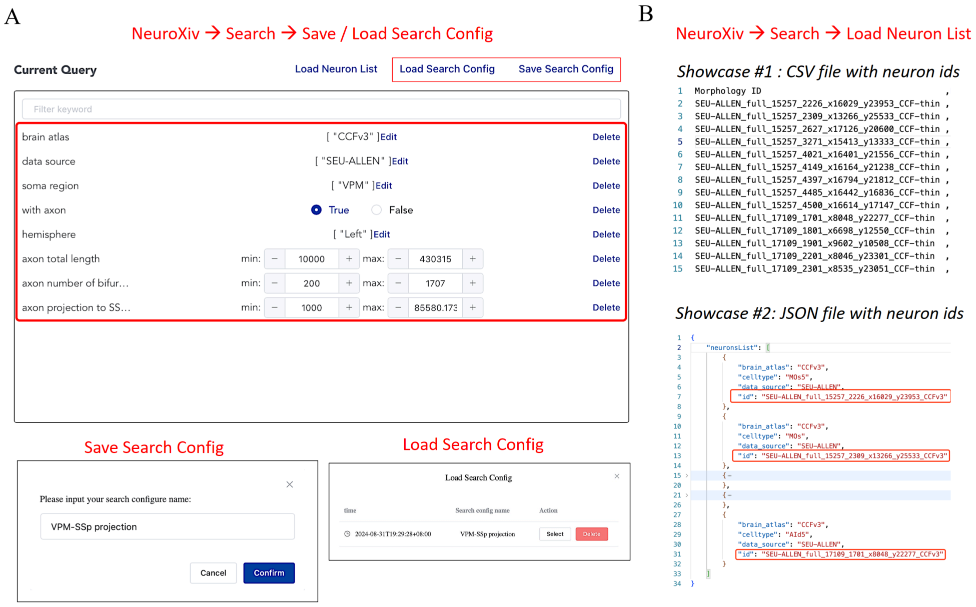


**Supplementary Fig. 4 | Additional search functions in NeuroXiv.** ***A****, search criteria can be saved or loaded.* ***B****, an illustration of how to utilize the database interface provided by NeuroXiv to view customized neuron data. Neuron data can be imported into NeuroXiv via CSV or JSON file.*

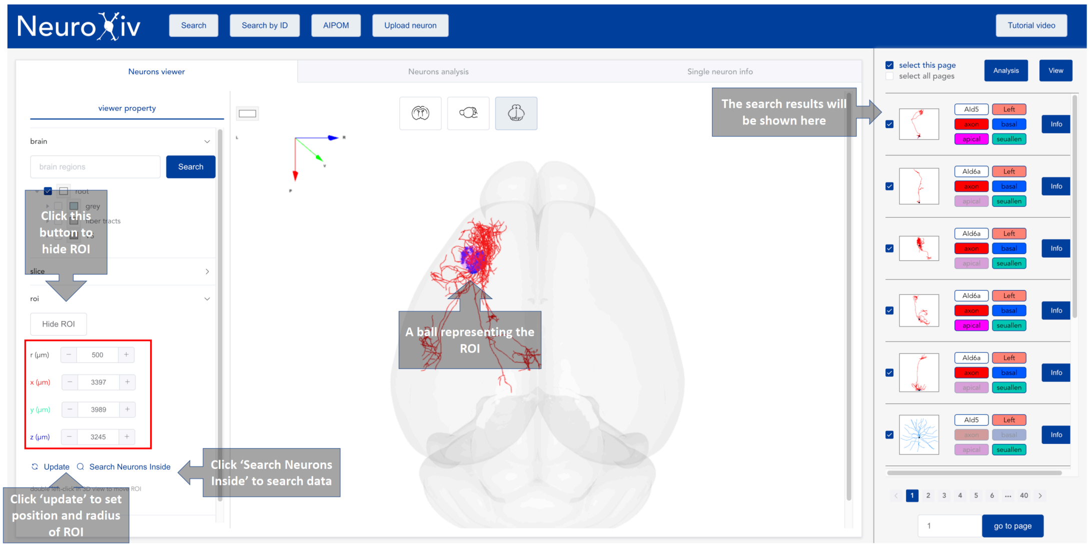


**Supplementary Fig. 5 | A screenshot demonstrating how to define a region of interest (ROI) in atlas to search for neurons.**

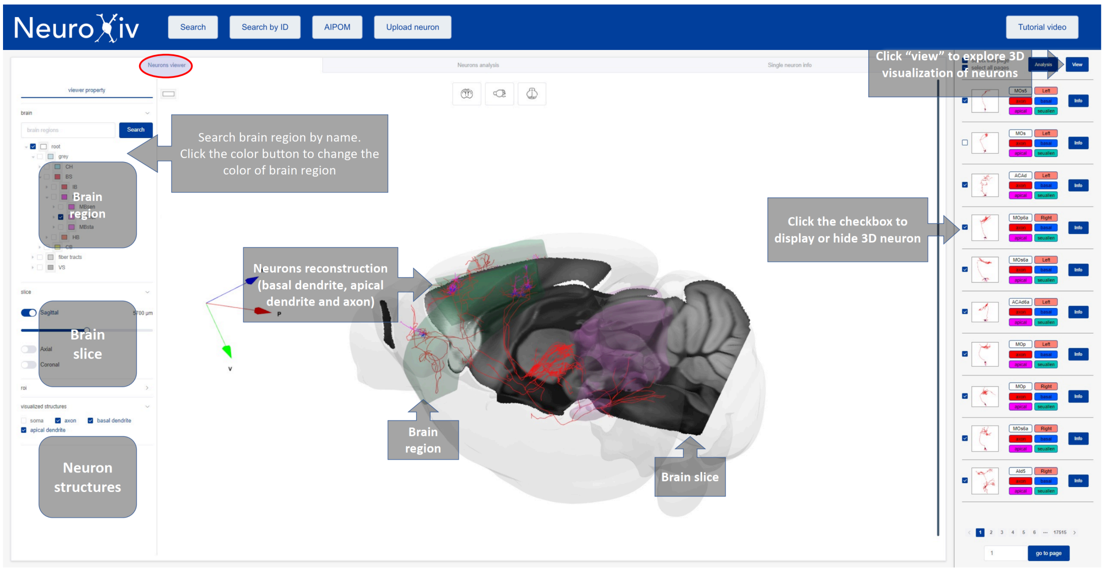


**Supplementary Fig. 6 | A screenshot demonstrating how to visualize neuron morphologies in atlas.**

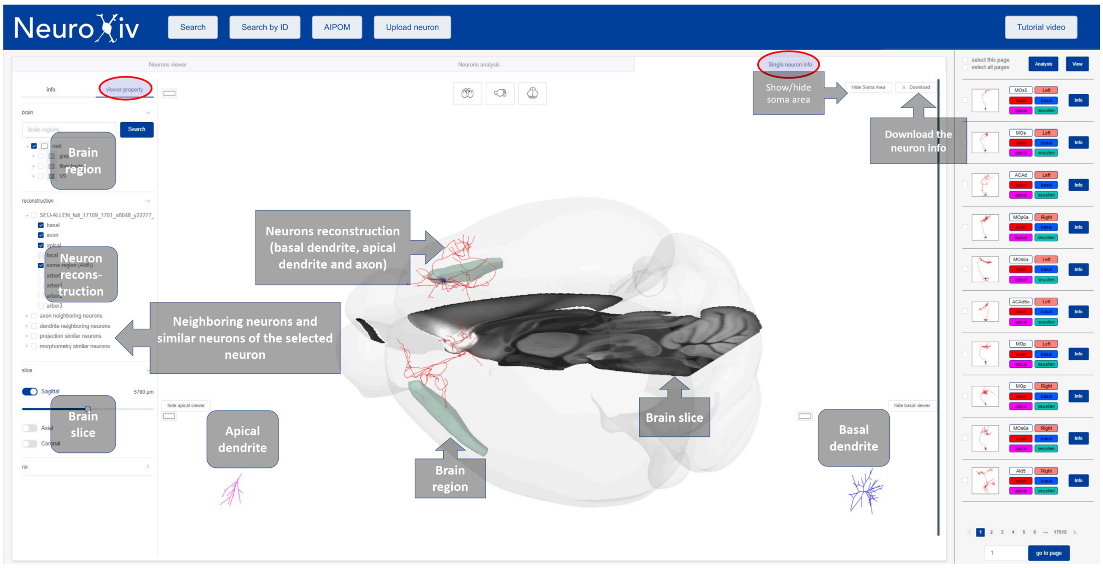


**Supplementary Fig. 7 | A screenshot demonstrating how to visualize the metadata and morphology of a neuron.**

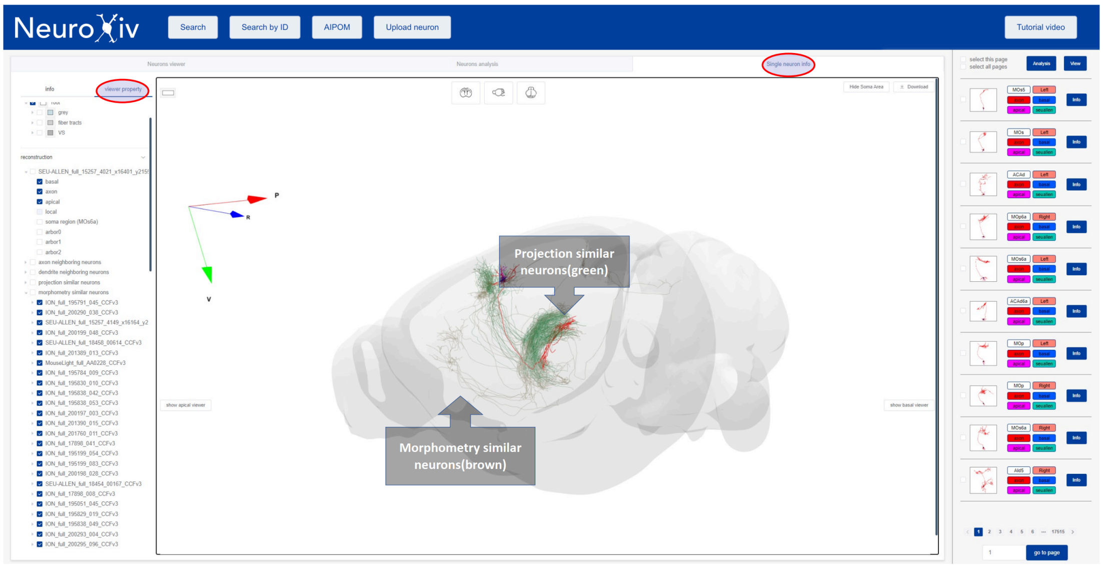


**Supplementary Fig. 8 | A screenshot demonstrating how to visualize similar neurons.**

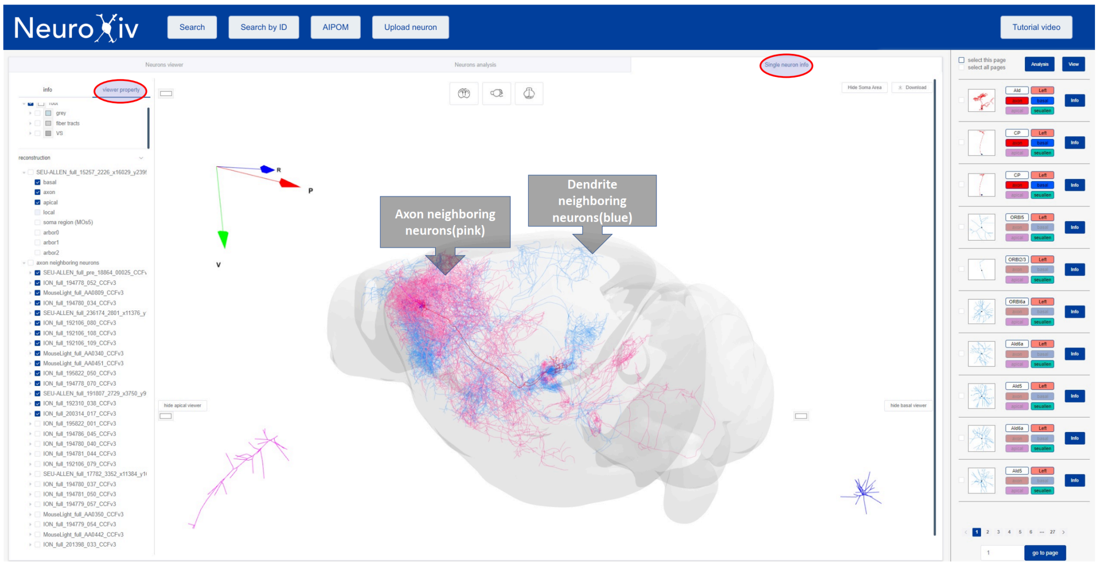


**Supplementary Fig. 9 | A screenshot demonstrating how to visualize neighboring neurons.**

### Supplementary Notes

#### Supplementary Note 1: Challenges in reusing the open available neuron morphology datasets

Neuron morphology plays a crucial role in classifying cell types and exploring functional connectivity within the brain (Luo, 2021; Zeng, 2022). Recent advancements in labeling (Aransay et al., 2015; Karube et al., 2004; Rotolo et al., 2008), imaging (Economo et al., 2016; Gong et al., 2016), and tracing (Bria et al., 2016; Jiang et al., 2022; Peng et al., 2017; Y. Wang et al., 2019) techniques have facilitated the accumulation of extensive neuron morphology datasets (Gao et al., 2022, 2023; Peng et al., 2021; Qiu et al., 2024; Winnubst et al., 2019) on a whole-brain scale. Numerous projects (Callaway et al., 2021; Hawrylycz et al., 2023; Manubens-Gil et al., 2023) have been initiated for specific research objectives, resulting in varied scales of neuron morphology datasets distributed across specific brain regions.

While dataset producers typically conduct comprehensive analyses and share raw neuron morphology data, integrating these datasets for comprehensive studies—such as result validation, secondary development, or cross-dataset comparisons—remains challenging (Martone, 2024; Wilkinson et al., 2016). Users are required to possess substantial domain expertise and advanced computational skills to manage the complexities of data collection, cleaning, standardization, registration, management, and analysis (Jiang et al., 2022; Y. Liu et al., 2023). Furthermore, substantial hardware resources are often necessary to meet the computational demands associated with processing large-scale neuron morphology datasets.

**Data Collection**

Data collection in neuron morphology research is a multifaceted challenge that requires researchers to locate relevant datasets based on their specific research goals. These datasets may be hosted across a variety of platforms, including private servers, general-purpose cloud storage services (e.g., AWS or Google Drive), version-controlled repositories (e.g., GitHub), and specialized repositories such as NeuroMorpho.org. Navigating these platforms requires understanding their capabilities, limitations, data accessibility, and metadata availability. Ensuring data integrity and completeness across these sources is crucial for downstream analysis. Each platform offers unique advantages, such as robust metadata support or community-curated datasets, but also presents challenges related to data versioning, consistency in updates, and interoperability with analytical tools. Effective data collection strategies require familiarity with these platforms, the ability to integrate datasets seamlessly, and rigorous data provenance and quality assurance protocols.

**Data Standardization**

Standardizing neuron morphology data poses challenges in converting data into universally supported formats, such as the SWC format (Mehta et al., 2023), essential for interoperability across platforms and tools. It also requires harmonizing classification and naming conventions for neuronal structures (e.g., soma, dendrite, axon) to ensure consistency in interpretation and analysis. Additionally, standardizing data coordinates within a common spatial framework (discussed in the Atlas Mapping section) is crucial for the accurate spatial alignment of neuronal reconstructions across different experiments and species.

Achieving effective standardization necessitates specialized tools for file conversion, data annotation, and quality control. These tools ensure rigorous validation and normalization, supporting reliable data analysis. Standardization enhances data usability, fosters collaboration, and facilitates meta-analyses in neuroscience. However, it requires detailed attention to both neuroanatomy and computational tools.

**Atlas Mapping**

Mapping neuron morphology data to a common coordinate framework, such as the Allen Brain Atlas's CCF (Q. Wang et al., 2020), enables spatial alignment and data integration across diverse sources. This process involves registering neuron reconstructions to standardized anatomical templates, allowing precise localization within brain regions and across different scales. The challenge arises from the existence of multiple brain atlases, each constructed using varied datasets and methodologies. Different atlases are tailored to specific neuron morphology datasets. For example, whole-brain imaging datasets generated using fMOST techniques (Gong et al., 2016; Zhong et al., 2021) align best with atlases like CCF-thin, constructed using mBrainAligner (Li et al., 2022; Qu et al., 2022), whereas atlases like CCF-ME in NeuroXiv platform use neuronal feature spaces to delineate finer brain regions.

Comprehensive data analysis requires not only ensuring data alignment within the same atlas but also selecting the most appropriate atlas that suits the specific dataset characteristics. This allows researchers to compare neuronal features and connectivity patterns across different atlases, thereby advancing our understanding of brain organization and function.

**Metadata Extraction**

Extracting metadata from neuron morphology datasets is complex due to the variability in the data. Metadata includes essential information such as soma region and contained structures, as well as features crucial for analysis, like morphological and projection strength data. However, metadata can vary across datasets, and often, it is not openly available. Extracting metadata requires specialized tools, as neuron morphology data involve multiple structures (e.g., axons, dendrites), each necessitating different feature sets for quantitative description. This often necessitates the use of multiple tools (Akram et al., 2018) or even the development of custom solutions (Y. Liu et al., 2023). Standardized metadata schemas, controlled vocabularies, and automated extraction tools are critical to ensuring consistency, enhancing data interoperability, and supporting reproducibility across neuroscience studies.

**Analysis and Mining**

Analyzing neuron morphology data involves sophisticated statistical and computational techniques to characterize morphological and projection diversity (Y. Liu et al., 2023; Peng et al., 2021), identify distinct neuron types (Gao et al., 2022, 2023), and study single-cell level circuits (Jiao et al., 2023; L. Liu et al., 2023). These analyses require robust pipelines that integrate various tools for data retrieval, visualization, filtering, and feature extraction. Techniques such as feature dimension reduction, clustering, and machine learning algorithms are essential for uncovering patterns within complex datasets. However, the challenges go beyond tool usage and pipeline design. Researchers must also synthesize diverse analytical results to extract meaningful insights and generate new hypotheses about brain organization and function.

Neuron morphology data aggregation presents a wide array of challenges, spanning technical and conceptual domains. Successfully addressing these challenges requires a deep understanding of the biological context as well as the computational tools needed to process, standardize, map, and analyze large-scale datasets. Moreover, the ability to visualize and interact with these data in three-dimensional spaces, such as the Common Coordinate Framework (CCF), adds another layer of complexity. By overcoming these challenges through meticulous data curation, robust standardization practices, and advanced analytical techniques, researchers can unlock new insights into brain structure and function, driving forward progress in neuroscience.

### Supplementary Note 2: Comparison with other platforms

NeuroXiv stands out in comparison to several well-established platforms, including the MouseLight Neuron Browser (MNB, http://ml-neuronbrowser.janelia.org), ION Mouse Projectome Atlas (IONMPA, https://mouse.digital-brain.cn/projectome), and NeuroMorpho (neuromorpho.org).

**Data Reusability**

The data on MNB is encoded in JSON files that describe neuron morphology and metadata, requiring users to develop custom programs for import. IONMPA provides data in the SWC file format, which is widely used but employs non-standard labeling conventions (e.g., axons labeled as type=0 instead of the standard type=2), necessitating label conversion. Additionally, some IONMPA data exhibits irregularities in tree structure definitions, such as non-uniquely connected neuron trees. NeuroMorpho aggregates data from over 900 labs globally, but users must apply specific search filters to find relevant datasets, and the data lacks standardization across different sources, often residing in disparate coordinate spaces rather than a unified framework like the Common Coordinate Framework (CCF).

In contrast, NeuroXiv provides data that is ready for immediate use. Our platform employs the standard SWC format for neuron morphology, with all data mapped to the same brain atlases, ensuring consistency. Moreover, NeuroXiv includes comprehensive metadata, simplifying downstream analysis and enabling users to seamlessly integrate and analyze data.

**Data Volume**

NeuroXiv hosts the largest collection of reconstructed neuron morphologies for the mouse brain, encompassing a wider variety of neuron types. We also offer multiple brain atlases to cater to different research needs, all accompanied by standardized metadata to facilitate comparative studies and enhance data utility.

**Data Retrieval**

A robust and user-friendly search tool is essential for effectively indexing data from large-scale databases, particularly given the complexity of tree-like neuron morphologies. Traditional data retrieval methods often require users to specify detailed search criteria, necessitating a high level of familiarity with both the data and its attributes. Additionally, mastering these tools typically involves a significant learning curve. Compared to other platforms, NeuroXiv excels in data retrieval capabilities. We provide visual filtering options, natural language-based search functionality, and advanced combinatorial search features based on morphological and projection characteristics, enabling more intuitive and precise data exploration.

**Data Analysis and Mining**

NeuroXiv offers the most extensive suite of tools for data analysis and mining. These include AI-based mining reports, tools for exploring neuron similarity, and resources for discovering neuron connectivity. These features empower users to gain deeper insights into the data, expanding the possibilities for scientific discovery.

**Open Access**

NeuroXiv is fully open access, requiring no user registration. Leveraging Amazon AWS cloud services, we provide global access to our data, metadata, and discovery datasets, making it easier for scientists worldwide to download and utilize these resources.

Overall, NeuroXiv offers significant utility through the standardization and centralization of neuron morphology data from various laboratories. Neuron morphologies are encoded in a unified file format and registered within a consistent coordinate space (brain atlases), ensuring data reusability and enabling cross-laboratory analyses. Corresponding metadata, including neuron types, morphological characteristics, projection features, and data correlations, are generated under different atlases and stored alongside all neuron data in a centralized cloud database, enhancing data retrievability. The AI-powered data report provides a robust tool for neuron analysis and data mining, and the platform is designed to easily accommodate additional types of analysis reports in the future. This scalability ensures that NeuroXiv can continue to evolve and meet the growing demands of the neuroscience community.
