## Extended Data Table 1 for "Beyond Static Brain Atlases: AI-Powered Open Databasing and Dynamic Mining of Brain-Wide Neuron Morphometry"

| Name | | NeuroXiv | MouseLight neuron browser | Digital Brain (ION) | NeuroMorpho |
|---|---|---|---|---|---|
| Website | | neuroxiv.org | mouselight.janelia.org | mouse.digital-brain.cn | neuromorpho.org |
| **Database** | number of neurons (mouse) | 177,186 | 1,227 | 16,788 | 137,457 (unknown completeness) |
| | number of cell types (anatomy) | 294 | 126 | 25 | 72 |
| | structural domains | soma, axon, (basal and apical) dendrite, arbor | soma, axon, dendrite | soma, axon, dendrite | soma, axon, (basal and apical) dendrite |
| | number of data sources | 3 | 1 | 1 | 13 |
| | data format and standard | standard SWC format | JSON file containing medata and neuron structures | SWC format file with non-standard and inconsistent structural domain identifiers | standard SWC format (no quality control) |
| | reference atlases | CCFv3 | CCFv2.5, CCFv3 | CCFv3 | Not exist one for all neurons |
| **Data query** | neuron browser | Yes (neuron thumbnail and metadata) | Yes (id based) | Yes (id based) | Yes (id based) |
| | by soma region | Yes | Yes | Yes | Yes |
| | by spatial coordinates | Yes | Yes | | |
| | by projected regions | Yes | Yes | Yes | |
| | by morphological features | Yes | | | Yes |
| | by nature language | Yes | | | |
| | by user-supplied neuron id (s) | Yes | | | |
| **Data visualization** | interactive visualization | Yes | Yes | Yes | 2D snapshot only |
| **Data analysis** | statistics | Yes | | Yes | Yes |
| | projection patterns | Yes | | Yes | |
| | morphological feature distribution | Yes | | | measures of one neuron |
| | AI report | Yes | | | |
| **Data mining** | morphology similar neurons | Yes | | | |
| | projection similar neurons | Yes | | | |
| | spatial neighboring neurons | Yes | | | |
| **Open access** | no authentication | Yes | Yes | | Yes |
| | (meta) data | Yes | Yes | morphology data only | Yes |
| | queried (meta) data | Yes | Yes (limited number : 20) | Yes (limited number : 500) | Yes |
| | discovery data | Yes | | only metadata of part of figures | |
| | processing tools / pipelines | Yes | | | Yes |
| | processing required before data reuse | | file format conversion and standardization, quality control, atlas mapping, metadata extraction | | |
