## Extended Data Table 2 for "Beyond Static Brain Atlases: AI-Powered Open Databasing and Dynamic Mining of Brain-Wide Neuron Morphometry"

| Metadata | Description |
| --- | --- |
| **ID** | Unique identifier in database |
| **data source** | Which institute is the data sourced from. |
| **soma region** | Soma region/cell type of the neuron. |
| **brain atlas** | The brain atlas that neurons mapped to. |
| **reconstruction method** | The types of data reconstruction methods. "semi-auto" indicates that the data was reconstructed through a combination of automated and manual processes, while "auto" indicates that the data was produced using a fully automated algorithm. |
| **with axon** | Whether the neuron has reconstructed axonal arbor |
| **with dendrite** | Whether the neuron has reconstructed (basal) dendritic arbor |
| **with apical dendrite** | Whether the neuron has reconstructed apical dendritic arbor |
| **hemisphere** | Which hemisphere of the brain atlas is the soma located in: "Left" or "Right" |

| | |
|---|---|
| **Center shift** | Center shift is defined as the distance between soma and the centroid of neuron which is computed by the weighted average of spatial position of all nodes in neuron reconstruction. Nodes are weighted by the distance to its parent node. Equal weight can be assigned to each node if a neuron reconstruction is resampled with fixed conterminous nodes distance. |
| **relative center shift** | Relative center shift is defined as the ratio of center shift and maximum Euclidean distance. |
| **average contraction** | Contraction is defined as the ratio of the Euclidean distance and the path distance of a branch. A branch always starts at soma/bifurcation and ends at next bifurcation/tip. Average contraction is the mean value of contractions of all branches. |
| **average bifurcation angle remote** | Given a bifurcation node A, it has two child branches. Each branch either lead to the next bifurcation or an ending tip. Assuming the nodes of the next bifurcation/ending tip is B_1 and B_2, the remote bifurcation angle is the angle of B_1 AB_2. The average bifurcation angle remote is the average remote angle of all bifurcations. |

| | |
|---|---|
| **average bifurcation angle local** | Given a bifurcation node A and its child nodes C_1 and C_2, the local bifurcation angle is the angle of C_1 AC_2. The average bifurcation angle local is the average local angle of all bifurcations. |
| **max branch order** | The maximum value of the order of branch. The order of the branches directly connect to soma is 0 and the order will increase by 1 after each bifurcation such that the number of bifurcations between a branch and the soma is its order. |
| **number of bifurcations** | The number of bifurcations in a neuron. As we use binary tree structure to represent neuron reconstruction, the bifurcation refers to a tree node (excluding soma) which has two child nodes. |
| **total length** | The sum of distance between two conterminous nodes. |
| **max euclidean distance** | The maximum value of Euclidean distance between node and soma. |
| **max path distance** | The maximum value of path distance between node and soma. |
| **average euclidean distance** | The average value of Euclidean distance between node and soma. |

| | |
|---|---|
| **25% euclidean distance** | The first quartile value of the Euclidean distance between node and soma. |
| **50% euclidean distance** | The median value of the Euclidean distance between node and soma. |
| **75% euclidean distance** | The third quartile value of the Euclidean distance between node and soma. |
| **average path distance** | The average value of path distance between node and soma. |
| **25% path distance** | The first quartile value of the path distance between node and soma. |
| **50% path distance** | The median value of the path distance between node and soma. |
| **75% path distance** | The third quartile value of the path distance between node and soma. |

| | |
|---|---|
| **2d density** | 2D Density=Pixels/Area Pixels is the number of pixels occupied by neuron arbors after projecting neuron reconstruction on the XY-plane. Specifically, we set node diameter and pixel size as 1um. To compute Pixels, neuron reconstruction is resized such that its spatial coordinate scale is 1um isotropic. The (x,y) coordinate of each node is rounded as integers and a set of distinct position vectors (x,y) is established. Pixels is obtained by counting the number of elements in the set. Area is estimated following the same method previously introduced. |
| **3d density** | 3D Density=Voxels/Volume Voxels is the number of voxels occupied by neuron arbors in 3 dimensional space. Specifically, we set node diameter and voxel size as 1um. To compute Voxels, neuron reconstruction is resized such that its spatial coordinate scale is 1um isotropic. The (x,y,z) coordinate of each node is rounded as integers and a set of distinct coordinate vectors (x,y,z) is established. Voxels is obtained by counting the number of elements in the set. Volume is estimated following the same method previously introduced. |

| | |
|---|---|
| **area** | Area was calculated by the area of polygon convex hull after projecting neuron on XY-plane. The convex hull was calculated by the 'scipy.spatial.ConvexHull' function using python package 'SciPy' (https://docs.scipy.org). Our measuring unit is um^2. |
| **volume** | Volume was calculated by the volume of convex hull in the 3 dimensional space. The convex hull was calculated by the 'scipy.spatial.ConvexHull' function using python package 'SciPy' (https://docs.scipy.org). Our measuring unit is um^3. |
| **width** | Width is computed by the difference between maximum and minimum node coordinate on y-axis. |
| **width 95ci** | Width 95%CI is computed by the difference between upper bound and lower bound of 95% confidence interval of node coordinate on y-axis. |
| **height** | Height is computed by the difference between maximum and minimum node coordinate on x-axis. |
| **height 95ci** | Height 95%CI is computed by the difference between upper bound and lower bound of 95% confidence interval of node coordinate on x-axis. |

| | |
|---|---|
| **depth** | Depth is computed by the difference between maximum and minimum node coordinate on z-axis. |
| **depth 95ci** | Depth 95%CI is computed by the difference between upper bound and lower bound of 95% confidence interval of node coordinate on z-axis. |
| **slimness** | The ratio of 'Width' and 'Height'. |
| **slimness 95ci** | The ratio of 'Width 95%CI' and 'Height 95%CI'. |
| **flatness** | The ratio of 'Height' and 'Depth'. |
| **flatness 95ci** | The ratio of 'Height 95%CI' and 'Depth 95%CI'. |
| **projection (relative)** | The ratio of the arbor length in this area and total arbor length |
| **projection (absolute)** | The arbor length in this area |
