## Extended Data Table 3 for "Beyond Static Brain Atlases: AI-Powered Open Databasing and Dynamic Mining of Brain-Wide Neuron Morphometry"

| query items | | server-side method (ms) | client-side method (ms) | Gain |
|---|---|---|---|---|
| query neuron types | search CA1 neurons | 16,599.4 | 1,687.2 | 9.8 |
| | search DG neurons | 14,286.5 | 1,569.9 | 9.1 |
| | search PL neurons | 19,531.8 | 1,975.4 | 9.9 |
| | search MOs neurons | 12,846.5 | 1,048.0 | 12.3 |
| | search CA3 neurons | 21,297.8 | 1,482.3 | 14.4 |
| | search Ald neurons | 24,380.3 | 3,622.0 | 6.7 |
| | search SUB neurons | 18,534.9 | 3,340.7 | 5.5 |
| | search ACAd neurons | 19,674.8 | 1,163.7 | 16.9 |
| | search VPM neurons | 13,016.1 | 1,435.2 | 9.1 |
| | search CP neurons | 12,664.5 | 2,642.9 | 4.8 |
| query neurons with particular structure | search CA1 neurons with axon | 24,787.2 | 1,630.7 | 15.2 |
| | search DG neurons with axon | 22,434.6 | 2,105.8 | 10.7 |
| | search PL neurons with axon | 24,884.9 | 1,138.6 | 21.9 |
| | search MOs neurons with axon | 27,150.4 | 1,178.4 | 23.0 |
| | search CA3 neurons with axon | 21,155.0 | 1,277.4 | 16.6 |
| | search Ald neurons with axon | 18,351.7 | 2,532.8 | 7.2 |
| | search SUB neurons with axon | 25,905.8 | 2,792.3 | 9.3 |
| | search ACAd neurons with axon | 17,380.2 | 3,053.6 | 5.7 |
| | search VPM neurons with axon | 37,543.3 | 2,326.7 | 16.1 |
| | search CP neurons with axon | 12,683.2 | 2,251.7 | 5.6 |
| query neurons with specific projection patterns | search CA1 neurons projecting to ACB | 15,865.6 | 2,568.2 | 6.2 |
| | search DG neurons projecting to CA3 | 33,939.7 | 4,821.4 | 7.0 |
| | search PL neurons projecting to CP | 31,327.7 | 2,796.8 | 11.2 |
| | search MOs neurons projecting to SSs | 59,691.9 | 2,008.7 | 29.7 |
| | search CA3 neurons projecting to LSr | 35,424.7 | 2,596.1 | 13.6 |
| | search Ald neurons projecting to MOs | 31,140.7 | 2,357.7 | 13.2 |
| | search SUB neurons projecting to MM | 24,500.8 | 2,155.4 | 11.4 |
| | search ACAd neurons projecting to CP | 49,261.0 | 2,418.1 | 20.4 |
| | search VPM neurons projecting to MOp | 26,355.9 | 2,323.7 | 11.3 |
| | search CP neurons projecting to SNr | 31,316.8 | 2,203.5 | 14.2 |
| Average | | 24,797.8 | 2,216.8 | 12.3 |
